# Frequency selectivity of persistent cortical oscillatory responses to auditory rhythmic stimulation

**DOI:** 10.1101/834226

**Authors:** Jacques Pesnot Lerousseau, Agnès Trébuchon, Benjamin Morillon, Daniele Schön

## Abstract

Cortical oscillations have been proposed to play a functional role in speech and music perception, attentional selection and working memory, via the mechanism of neural entrainment. One of the most compelling arguments for neural entrainment is that its modulatory effect on ongoing oscillations outlasts rhythmic stimulation. We tested the existence of this phenomenon by studying cortical neural oscillations during and after presentation of melodic stimuli in a passive perception paradigm. Melodies were composed of ∼60 and ∼80 Hz tones embedded in a 2.5 Hz stream. Using intracranial and surface recordings in humans, we reveal consistent neural response properties throughout the cortex, well beyond the auditory regions. Persistent oscillatory activity in the high-gamma band was observed in response to the tones. By contrast, in response to the 2.5 Hz stream, no persistent activity in any frequency band was observed. We further show that our data are well-captured by a model of damped harmonic oscillator and can be classified into three classes of neural dynamics, with distinct damping properties and eigenfrequencies. This model provides a mechanistic and quantitative explanation of the frequency selectivity of auditory neural entrainment in the human cortex.

**Significance statement:** It has been proposed that the functional role of cortical oscillations is subtended by a mechanism of entrainment, the synchronisation in phase or amplitude of neural oscillations to a periodic stimulation. We tested whether the modulatory effect on ongoing oscillations outlasts the rhythmic stimulation, a phenomenon considered to be one of the most compelling arguments for entrainment. Using intracranial and surface recordings of human listening to rhythmic auditory stimuli, we reveal consistent oscillatory responses throughout the cortex, with persistent activity of high-gamma oscillations. On the contrary, neural oscillations do not outlast low-frequency acoustic dynamics. We interpret our results as reflecting harmonic oscillator properties - a model ubiquitous in physics but rarely used in neuroscience.

## Introduction

Cognitive neuroscience aims to determine the nature of the basic computations underlying cognition and how they are implemented ^1–3^. Neural oscillations are an emergent property of a population of interacting neurons that can be described by few phenomenological parameters, such as phase, amplitude and frequency ^4–6^. According to recent theories, neural oscillatory activity plays a crucial role in the implementation of elemental computations ^7–9^. Indeed, oscillations have the powerful property of indexing algorithms, such as hierarchical parsing ^10^, chunking ^11^ and clocking ^12^, while being easy to describe mechanistically. From a cognitive neuroscience angle, they constitute an important interface between algorithmic and implementational levels of analysis ^13^.

Rhythmic stimulation, either sensory or electrical, can be used to modulate neural oscillatory activity, to reveal or optimize brain functions ^14, 15^. Such modulation is believed to be realised through neural entrainment, defined here as the synchronisation in phase or amplitude of neural oscillatory activity to a periodic stimulation ^16, 17^. At the algorithmic level, entrainment phenomena have been proposed to account for segmentation of speech ^18–23^, attentional selection during visual or auditory perception ^24–29^, perception of the musical beat ^30–32^ and auditory working memory performance ^33^. At the implementational level, however, entrainment can occur only if the stimulation is applied at a frequency close to an eigenfrequency of the targeted cell assembly. Thus, depending on whether it is applied at or away from a network’s eigenfrequency and whether neural oscillations are self-sustained or not, a rhythmic stimulation will either induce oscillatory entrainment, oscillatory resonance or a superposition of transient event-related potentials ^34, 35^. This distinction is at the heart of a vibrant debate concerning the nature of neurophysiological responses such as steady-state (SSR) ^29, 36^, frequency-following (FFR) ^37–39^ or envelope-following (EFR) responses ^40–44^.

One of the most compelling arguments for entrainment is that its modulatory effect on ongoing oscillations outlasts rhythmic stimulation ^45^. In dynamical systems approaches, it corresponds to the fundamental property of underdamping (damping ratio *ζ* < 1), *i.e.* the ability for a system to maintain a long-lasting oscillation, echo or reverberation, whose amplitude exponentially decreases toward baseline after the stimulation ends ^44, 46, 47^. Strikingly, this property has never been systematically investigated despite being considered as a possible mechanism underlying temporal predictions in multiple cognitive theories, in particular at low (< 10 Hz) frequencies: dynamic attending theory ^30^, multisensory integration ^48, 49^, timing ^12^, or interpersonal interaction ^50^. In the auditory domain especially, where the temporal structure of sound streams is highly informational, either for speech comprehension ^10, 51, 52^ or musical-beat perception ^30, 31, 36, 41^, underdamping is believed to be the property that allows the brain to anticipate the sounds.

To investigate the damping properties of cortical oscillations during auditory rhythmic stimulation, we recorded whole-brain cortical neurophysiological activity with either stereotactic electroencephalography (sEEG) on epileptic patients implanted for clinical evaluation, or magnetoencephalography (MEG) on healthy participants. While sEEG provides the best spatiotemporal resolution and signal-to-noise ratio of human cortical recordings ^53^, optimizing our chances to detect long-lasting oscillations, it does not provide a full cortical coverage, which is afforded by the complementary MEG recordings ^54^. Participants passively and repetitively listened to a 6 s auditory stream composed of high frequency tones (∼80 or ∼60 Hz) presented at a rate of 2.5 Hz. Periods of silence separated both successive tones and streams, to allow investigating the damping properties of neural oscillatory responses. We explored the occurrence of cortical neural oscillations during and after presentation of melodic stimuli at both tones (high, ∼80/60 Hz) and stream (low, 2.5 Hz) frequencies.

Based on the auditory literature, we hypothesized that neural underdamping would be selective to low-frequency stimulation and would mainly occur in auditory, but also motor cortical regions ^31, 55^. Hence, while each tone would evoke a high-gamma (∼60/80 Hz) neural response that would end at stimulus offset, the underlying rhythm would entrain delta (2.5 Hz) neural oscillations, characterized with underdamping, *i.e.* long-lasting activity. In addition to these phase-locked responses, we expected to observe an induced response in the beta range (15-30 Hz), whose amplitude would be entrained at the stream rate ^31, 56, 57^. In the following, we thus investigated all possible forms of phase and amplitude synchronisation, at all frequencies and in all cortical regions, in response to our composite auditory rhythmic stimulation. Hence, we investigated:

1. Whether underdamping occurs at all after passive auditory rhythmic stimulation.
2. If so, whether it depends on the frequency of stimulation.
3. Whether it is region and/or frequency specific.

## Results

### High-frequency acoustic modulations (∼60/80 Hz) induce long-lasting neural frequency-following responses

We first investigated the FFR, *i.e.* the neural evoked response at the fundamental frequency of the tones. The main goal of this analysis was to identify sEEG/MEG sources for which a FFR was observed and then to estimate whether this activity outlasted the duration of the stimulus. For this purpose, we developed a method that we refer to as Align-Bin-and-Count (ABC, see Methods, Fig. Supp. 1), in which we treat each oscillatory cycle as a bin of activity and estimate how many consecutive active bins are present in the neural signal. This method accounts for differences in power between sources, lag between stimulus and neural response and spurious oscillations arising from filtering smearing (see Methods). This allows us to accurately estimate the duration of each significant neural response and infer whether it outlasts stimulus duration or not.

In order to study the FFR, we applied ABC to the envelope of narrow-band filtered signals (83 Hz or 62 Hz). Importantly, we confirmed the validity of this method by applying the filter and ABC to the sound stimuli themselves, which yielded an accurate estimate of their number of cycles (see Fig. Supp. 2). Long-lasting activity duration could therefore not arise from spurious filtering smearing ^58^.

Analyses of sEEG recordings first reveal that FFR are present in a wide cortical network, extending well beyond auditory cortex (13 % of all recorded sites for 83 Hz, 21 % for 62 Hz, see Figure 1J-K; white and coloured dots, for consistency across participants see Fig. Supp. 3). This network comprises auditory regions (Heschl gyrus and planum temporale bilaterally, extending to superior and middle temporal gyri) and also comprises pre-motor and motor regions (bilateral precentral gyrus and right premotor cortex) as well as associative regions involved in higher-level auditory processing (bilateral inferior parietal lobule and left inferior frontal gyrus). The FFR persists beyond stimulus duration in a substantial proportion of the responding sEEG sources (118/424 corresponding to 28 % at 83 Hz and 232/669 corresponding to 35 % at 62 Hz). The number of cycles of oscillatory activity can exceed that of the stimulus by up to 11 cycles at 83 Hz and 8 cycles at 62 Hz (∼180% of stimulus duration, see Figure 1H-I). The anatomical location of sEEG sources whose activity outlasts stimulus duration are also comprised within the entire dual-stream auditory cortical processing pathway ^59^ (see Figure 1J-K): in the antero-ventral stream, from auditory to inferior frontal cortex (auditory cortex, medial part of the superior temporal gyrus, middle temporal gyrus and left pars triangularis of the inferior frontal gyrus) and in the postero-dorsal stream, from auditory to motor cortex (posterior part of the superior temporal gyrus, inferior parietal lobule, precentral gyrus, and right premotor cortex).

**Figure 1.**
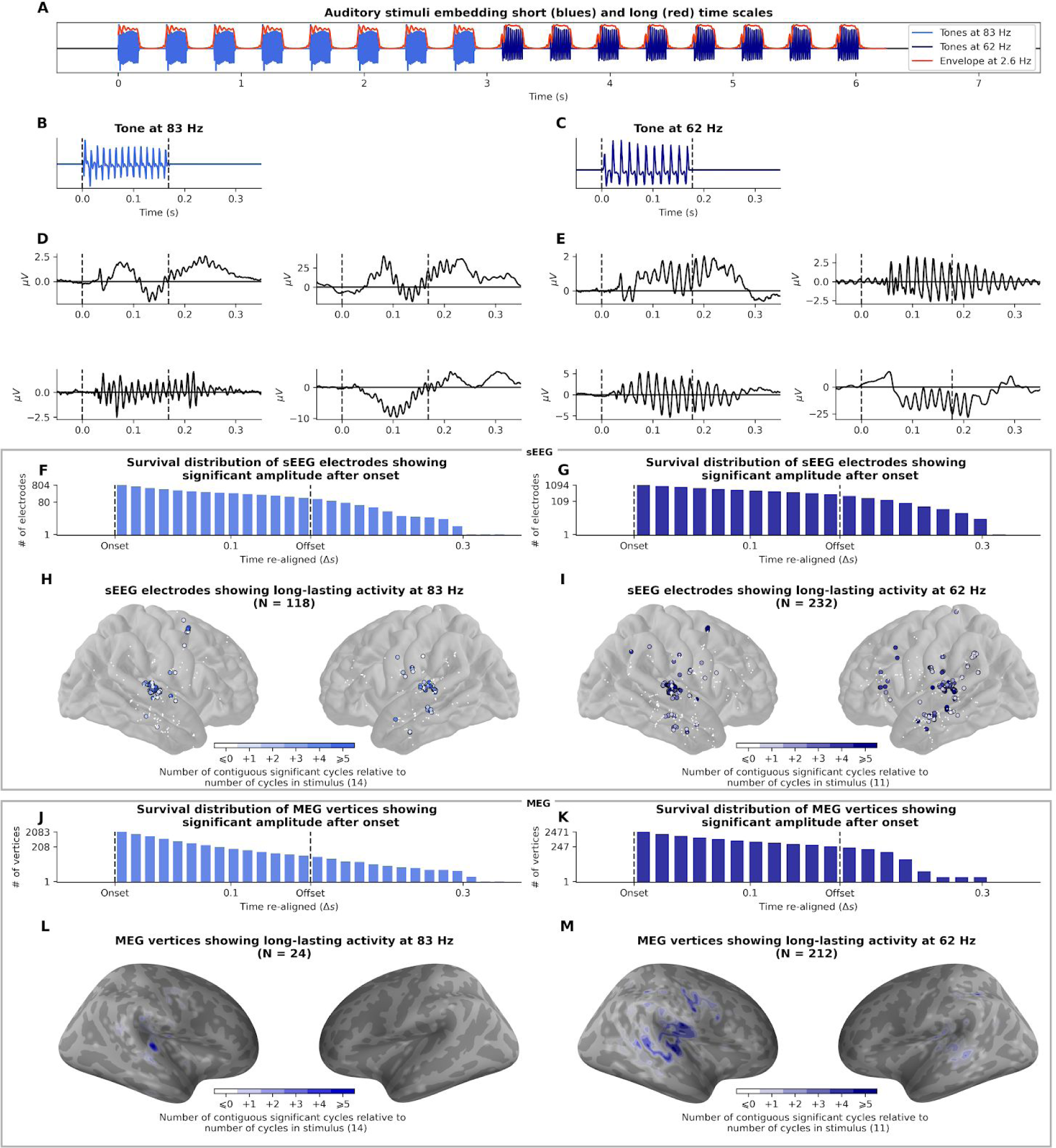
Estimation of frequency-following response duration (FFR, 83 and 62 Hz). **A.** Auditory stimulus of 8 tones with high frequency carriers at 83 Hz followed by 8 tones at 62 Hz presented at a 2.5 Hz rate (envelope). **B-C.** Waveform of the 83 and 62 Hz tones, made-up of 14 and 11 oscillatory cycles respectively. **D-E.** Evoked unfiltered response of 8 example sEEG channels, from different participants and different brain regions (Heschl’s gyrus, motor cortex and superior temporal sulcus). **F-G.** Survival distribution of sEEG channels with consecutive significant activity after onset. 118 and 232 channels outlast the 0.170 s duration of the 83 and 62 Hz stimuli, respectively. **H-I.** Localization of sEEG channels showing a FFR during the stimulus. Channels with activity outlasting stimulus duration are circled. Color intensity indicates the number of cycles outlasting stimulus duration. **J-K.** Survival distribution of MEG vertices activity showing consecutive significant activity after onset. **L-M.** Localization of MEG vertices showing a FFR. Color intensity indicates the number of cycles outlasting stimulus duration.

MEG data substantiates the sEEG results, revealing FFR in superior and inferior temporal gyri, precentral gyrus and supramarginal gyrus. Again, the FFR persists beyond stimulus duration (24/395 corresponding to 6 % at 83 Hz and 212/858 corresponding to 25 % at 62 Hz). The number of cycles of oscillatory activity can exceed that of the stimulus by up to 12 cycles at 83 Hz and 8 cycles at 62 Hz (∼180% of stimulus duration, see Figure 1L-M, Fig. Supp. 4). Similarly, the localisation of the MEG sources whose activity outlasts stimulus duration comprises regions of the auditory network (superior temporal gyrus, inferior parietal lobule and right precentral gyrus).

### Low-frequency acoustic modulations (2.5 Hz) do not induce long-lasting neural envelope-following responses

We used the same approach to investigate the neural response to the low-frequency amplitude modulations of the auditory stream, known as envelope-following response (EFR). Each tone of sequence was separated by a fixed inter-onset interval of 390 ms, resulting in a 2.5 Hz amplitude-modulated temporal envelope (see Figure 2B). We applied ABC to the envelope of the 2.5 Hz narrow-band filtered signals (see Figure 2C).

**Figure 2.**
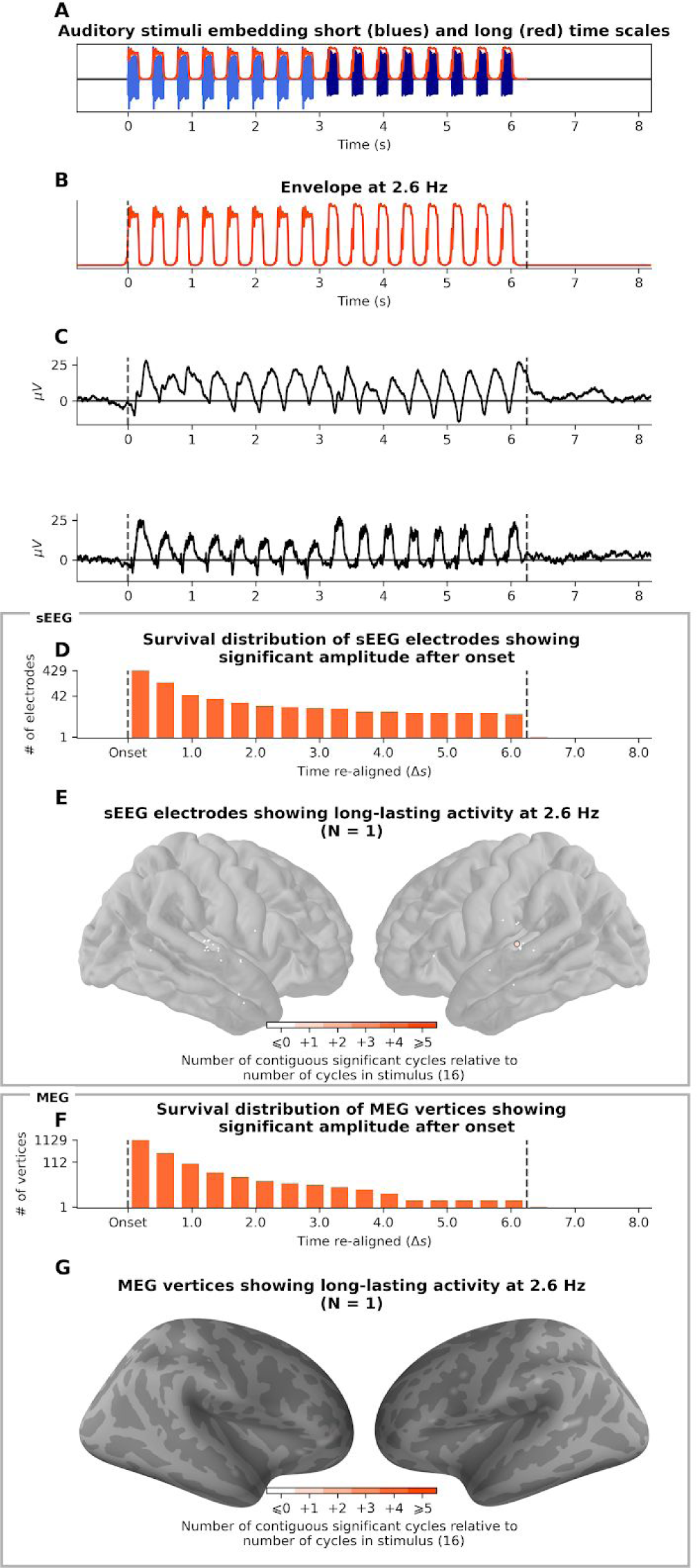
Estimation of envelope-following response duration (EFR, 2.5 Hz). Auditory stimulus. **B.** Waveform of the envelope fluctuation at 2.5 Hz. **C.** Evoked unfiltered response of 2 example sEEG channels, from different participants, both in Heschl’s gyrus. **D.** Survival distribution of sEEG channels with consecutive significant activity after onset. Channels whose activity lasts more than 16 cycles outlast the duration of the stimulus (N = 1). **E.** Localization of sEEG channels having an EFR. The channel with long lasting activity is circled. Color intensity indicates the number of cycles of post-stimulus activity (1). **F.** Survival distribution of MEG vertices activity showing consecutive significant activity after onset. **G.** Localization of MEG vertices that have an EFR. Color intensity indicates the number of cycles of post-stimulus activity.

SEEG recordings analyses reveal that, similarly to FFR, a wide cortical network presents an EFR (see Figure 2F). It comprises auditory regions (bilateral Heschl gyrus, planum temporale, and middle temporal gyrus), motor regions (bilateral precentral gyrus) as well as associative regions (left inferior frontal gyrus). However, unlike FFR, no long lasting EFR were observed. Only one (/40 responsive) sEEG source has a response that outlasts stimulus offset, and only by a single cycle (see Figure 2F). Similarly, MEG results did not reveal outlasting EFR (see Figure 2H, Fig. Supp. 5), except for one source (/32) exceeding by a single cycle the stimulus duration. Thus, low-frequency acoustic modulations do not induce outlasting ERF.

The two previous analyses investigated the N-to-N relationship between the stimulus frequency and the brain oscillatory activity: a frequency-specific neural oscillation (62 Hz, 83 Hz, or 2.5 Hz) in response to a frequency-specific oscillating stimulus. However, nesting -*e.g.* cross-frequency phase-phase or phase-amplitude coupling-phenomena exist and play an important role in shaping brain dynamics ^60–63^. We therefore broadened our analysis to all neural activity phase-locked to the stimulus. These N-to-M components were revealed by a time-frequency decomposition, using the inter-trial phase coherence (ITPC), a measure that quantifies the amount of phase consistency across trials for each frequency and time bin. A high ITPC is indicative of strong phase-locking to the stimulus for a particular frequency and a particular time point.

In order to investigate whether low-frequency (2.5 Hz) acoustic stimulation induces any long lasting ITPC, we thus decomposed the neural activity into five frequency bands of interest, defined based on the response of the auditory cortex. We estimated both the power time-frequency plane, quantifying the trial-averaged response amplitude of each frequency in time, and the ITPC plane. We then computed the mean and variance across time of both power and ITPC of each frequency during the presentation of the stimulus (see Figure 3, Fig. Supp. 6 for MEG). This resulted in five spectral distributions, revealing clear and recurring spectral peaks at which neural activity was modulated during stimulus presentation, in the delta (2-3.5 Hz), theta (4-7 Hz), alpha (8-11 Hz), beta (12-22 Hz) and gamma bands (50-110 Hz). In particular, we chose to investigate the beta band (12-22 Hz), as existing literature reports the implication of beta activity in the encoding of the temporal structure of acoustic stimuli ^31, 56, 57^.

**Figure 3.**
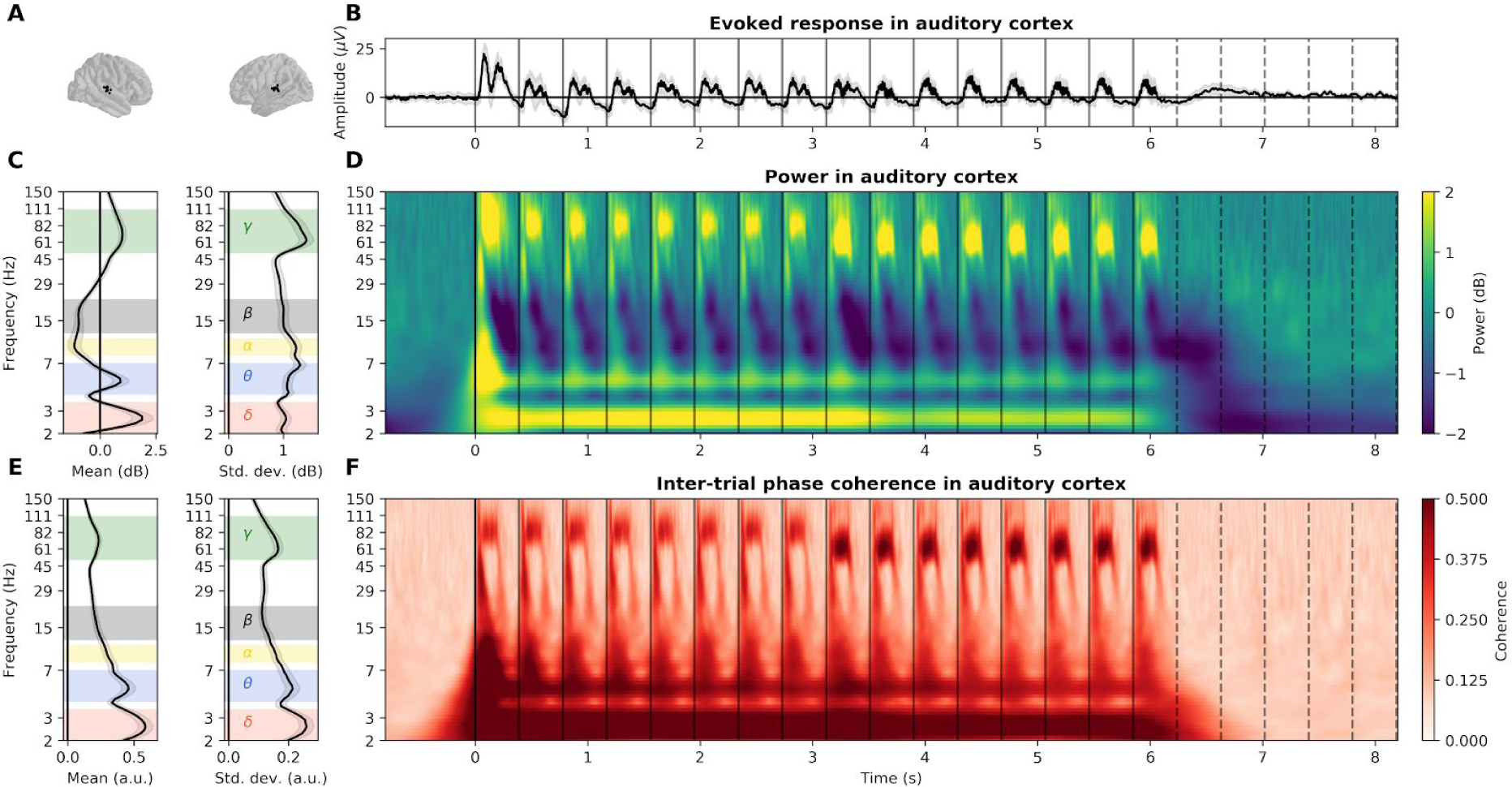
Detailed envelope-following response in auditory cortex recorded with sEEG. **A.** Localization of the sEEG channels located in bilateral auditory cortices. **B.** Evoked response, averaged across channels. Grey shaded area represents the standard error of the mean (s.e.m.) across sources. Vertical plain and dotted lines respectively indicate the onset of each tone and their theoretical continuation in the silence. **C**. (Left) Average and (right) standard deviation of power over time during stimulus presentation (0 - 6.2 s). Selected frequency bands are indicated by colored shaded areas (δ delta: 2-3.5 Hz, θ theta: 4-7 Hz, α alpha: 8-11 Hz, β beta: 12-22 Hz, γ gamma: 50-110 Hz). **D.** Power, averaged across channels, in dB relative to baseline. **E.** (Left) Average and (right) standard deviation of inter-trial phase coherence (ITPC) over time during the presentation of the stimulus. **F.** Inter-trial phase coherence (ITPC), averaged across channels. Note the presence of the FFR first at 80 and then at 60 Hz both in power and ITPC reflecting the change of tone fundamental frequency at ∼3 s.

We then applied the ABC procedure on all sEEG/MEG sources and all frequency bands ITPC to identify active sources and estimate their activity duration. SEEG and MEG recordings consistently reveal the spatial extent of the neural response for each defined frequency band (see Fig. Supp. 7; white and colored areas). While stimulation induces a large-scale evoked response -encompassing auditory, motor and associative regions-in the delta and theta bands, alpha, beta and gamma responses are more focal, in the bilateral superior temporal gyrus, and -for alpha and gamma responses-also in the precentral gyrus. Critically, none of these evoked responses outlasts significantly the stimulus duration. In the delta band only, a tiny proportion of channels (1% in sEEG, 6/531, <0.1% in MEG, 4/5124) outlasts the stimulus by one cycle.

The neural response can be split into phase-locked activity, revealed by ITPC, and induced activity, revealed by power averaging across trials ^64^. If the low frequency auditory modulation does not probe a long lasting phase-locked response, effects could still be present in the induced activity. For completeness, we therefore tested this hypothesis. We used machine learning to train a model (temporal response function ^65^, TRF) to decode the stimulus with a linear combination of induced power in all sources and all frequencies delayed in time. We reasoned as follows: if power is modulated by the envelope of the auditory stimulus, then these fluctuations are informative about the stimulus. We should therefore be able to train a TRF model to decode the envelope based on power data. If power continues to exhibit these fluctuations during the silence period directly following the offset of the stimulus, we should still be able to decode the envelope oscillations (see Methods). In sEEG (see Fig. Supp. 8A-B), the model performs remarkably well, predicting the stimulus envelope in the trained set with a coefficient of determination R^2^ of 0.97 (z = 22, p < 0.001 compared to surrogate distribution), and with an R^2^ of 0.71 in the testing set (z = 16, p < 0.001). When applied on the power directly following stimulus offset, the TRF performances drop to chance, with an R^2^ of -0.03 (z = -0.09, p = 0.57). Similar results were found in MEG (see Fig. Supp. 8C-D), where the model predicts the stimulus envelope in the trained set with a coefficient of determination R^2^ of 0.97 (z = 18, p < 0.001 compared to surrogate distribution), and with an R^2^ of 0.40 in the testing set (z = 7.4, p < 0.001). When applied on the power directly following stimulus offset, performance drops to chance, with an R^2^ of 0.03 (z = 0.43, p = 0.19). Thus, even when combining activity from all frequencies and sources, we are unable to detect information linked to the 2.5 Hz envelope modulation.

### A model of damped harmonic oscillator captures the dissociation between high and low frequency responses

Oscillatory phenomena are common in nature. The damped harmonic oscillator, used to describe very different systems, *e.g.* spring/mass systems, pendulums, torques and electrical circuits, is the standard model in physics. Despite being simple, powerful and well-suited to the description of neural mass dynamics ^66, 67^, this model has received little attention in the cognitive neuroscience community. By analogy to a spring/mass system (see Figure 4A), it is described by the canonical differential equation for linear oscillation :

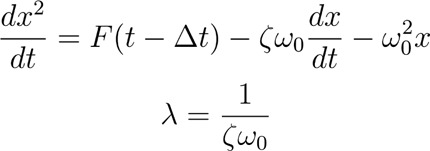

where *x* is the amplitude of the neural activity, *ζ* is the damping ratio, F is the stimulus amplitude, *Δt* is a time delay to account for transmission delays in the peripheral auditory system and 2*πω*_0_ is the eigenfrequency of the system. The damping ratio *ζ* is a key latent variable, as it constrains the activity of the system after the end of the stimulation. Over-damped systems (*ζ* > 1) show no oscillation after the end of the stimulation, whereas underdamped systems (*ζ* < 1) show oscillatory behavior with an amplitude decaying at an exponential decay of time constant *λ*. For example, a system with *ζ* = 0.1 and eigenfrequency 2*πω*_0_ = 1 Hz will take *λ*ln2 ≈ 7 s to return to half of its activity, i.e. ∼ 7 cycles after the end of the stimulation. Note that this goes beyond the debate on “self-sustained oscillators versus superposition of transient event-related potentials” insofar, as damping is a general property encompassing both interpretations. From a functional point of view, underdamping is the property that matters. The damped harmonic oscillator is thus a good model to clarify and quantify this question.

**Figure 4.**
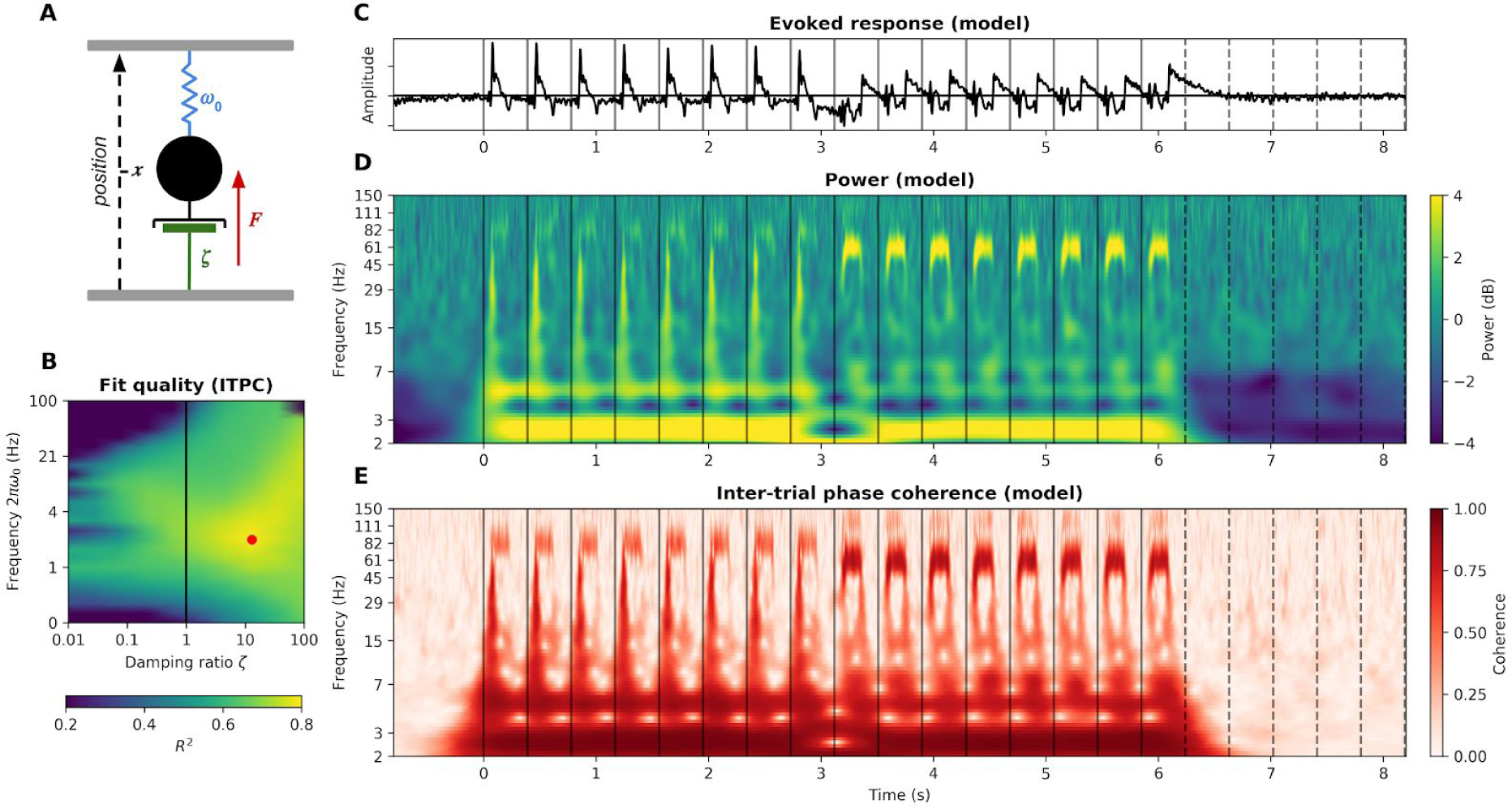
The auditory cortex as a damped harmonic oscillator. **A.** By analogy to a spring/mass system, the model is composed of the trajectory of a center of mass (neural amplitude, in black), a spring (a force pushing the system toward a fixed point, or baseline, in blue), a damper (dissipation of energy, phase dispersion, negative feedback, in green) and a driving force (the auditory stimulus amplitude, in red). **B.** Fit quality (R^2^) of the harmonic oscillator to the auditory cortex ITPC as a function of model parameters. Models with *ζ* < 1 are underdamped, model with *ζ* > 1 are overdamped. The best fitting model, highlighted in red (ζ = 13, 2πω_0_ = 2.1 Hz, Δ*t* = 40 ms) explains 78% of the ITPC variance. **C.** Evoked activity of the best fitting model. **D.** Power of the best fitting model. **E.** ITPC of the best fitting model.

The three free parameters of this model (*ζ*, 2*πω*_0_ and *Δt*) were fitted on the average power and ITPC time-frequency responses of the auditory cortex (see Methods). The model performs remarkably well, explaining 38% of the power variance and 78% of the ITPC variance. We next focussed on ITPC data. Although linear and simple, the model captures key aspects of the auditory cortex response (Figure 4D-E) : clear frequency following responses at 83 and 62 Hz, stronger responses to 62 than 83 Hz tones, onset and offset responses to each tone in all frequencies, strong phase coherence at 2.5 Hz as well as harmonics at 5 and 10 Hz. The best fitting model has parameters *ζ* = 13, 2*πω*_0_ = 2.1 Hz and *Δt* = 40 ms (Figure 4B). Its damping ratio *ς* >> 1 indicates strong overdamping, *i.e.* absence of oscillatory dynamics after the end of the stimulation, thus reproducing our previous results.

In order to differentiate neural populations with different dynamics, we fitted the harmonic oscillator model to each sEEG channel (see Figure 5, Fig. Supp. 9). After thresholding unexplained data (R^2^ < 5%, 257 channels survived), a clustering algorithm (k-means, optimal silhouette index at k=3, see Methods) yielded three clusters. Each cluster consists of a set of parameters and an associated topography. The first two clusters have a relatively low eigenfrequency 2*πω*_0_ (0.73 Hz [1^st^ decile: 0.43, 9^th^ decile: 2.1] and 2.1 Hz [1.3, 7.5]). Importantly and confirming our previous analyses, these two classes have a high damping ratio *ζ* (1.6 [1.0, 13] and 4.6 [0.6, 100]). Again replicating the model-free analyses for ITPC in the delta and theta band, the low frequency clusters comprise bilateral auditory regions, but also associative regions situated along the two auditory pathways (superior and medial temporal gyrus, precentral gyrus and bilateral inferior frontal gyrus). Conversely, the third cluster has radically different dynamics: a relatively high eigenfrequency 2*πω*_0_ (60 Hz [27, 100]) and low damping ratio *ζ* (0.08 [0.03, 0.6]). These parameters indicate an exponential decay of the amplitude of the oscillation after the end of the stimulation of time constant *λ* = 0.9 s [0.12, 2.2]. The sEEG channels constituting this cluster are located in auditory cortices, precentral gyrus, medial temporal gyrus and right inferior frontal gyrus. Overall, these three clusters confirm the dissociation between high and low frequency damping properties: the low frequency clusters (0.7 and 2 Hz) show overdamping, whereas the high frequency cluster (60 Hz) shows strong underdamping.

**Figure 5.**
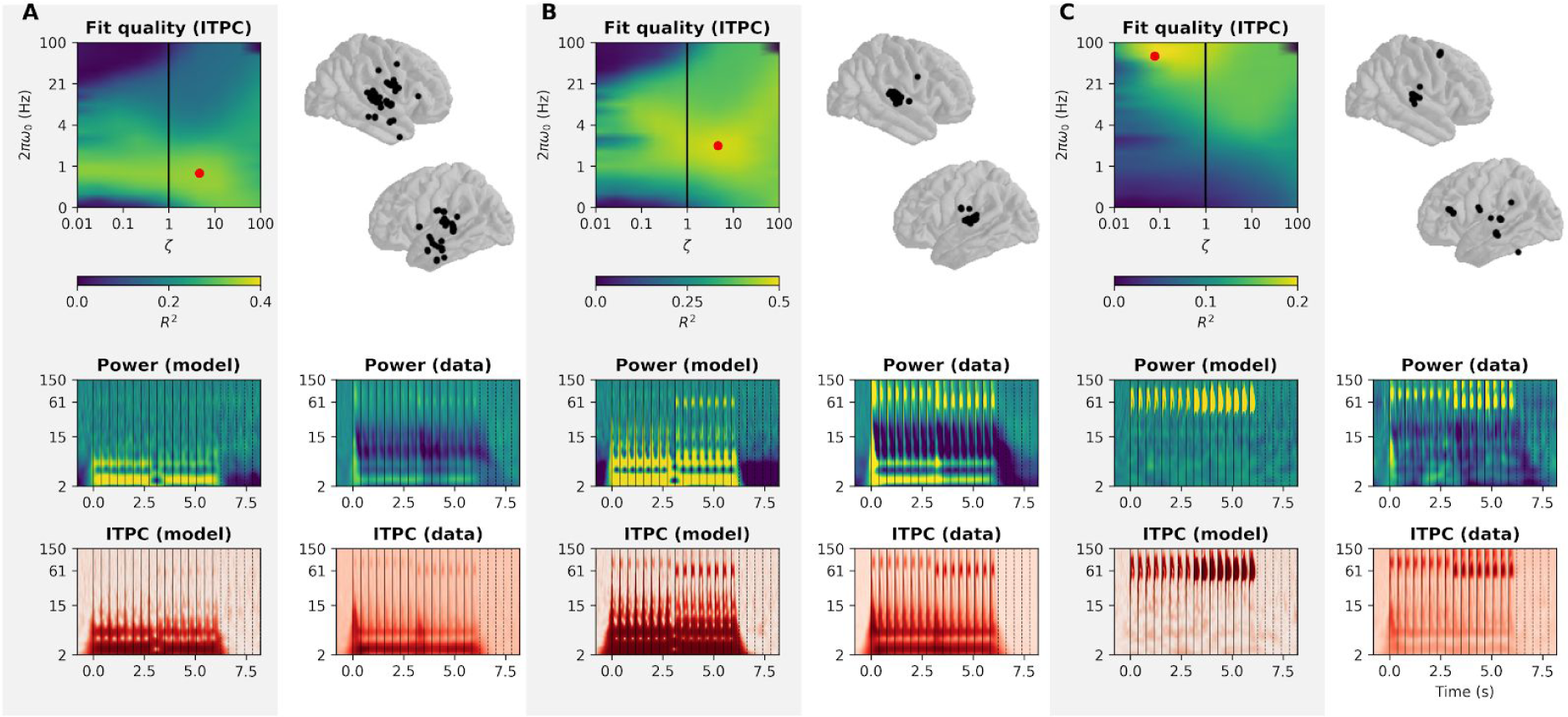
Clustering of electrodes based on best fitting parameters. **A.** Cluster #1. (Left) Average fit quality (R^2^) as a function of model parameters. The best fitting model is highlighted in red. (Right) Topography of the sEEG channels included in the cluster. (Down, left) Power and ITPC of the best fitting model for this cluster. (Down, right) Average power and ITPC of the sEEG channels included in the cluster. **B.** Cluster # 2. **C.** Cluster # 3.

## Discussion

During passive listening of a rhythmic stream of tones, the human brain exhibits two types of neural responses. First, a high-gamma oscillatory response, phase-locked to the fundamental frequency of the tones, that persists up to ten cycles after stimulus offset. Second, a complex set of responses encompassing all frequencies, evoked or induced by tones onset, offset, and the low-frequency acoustic rhythm, that does not persist long after stimulus offset (≤1 oscillatory cycle). These two responses are well-captured by three classes of damped harmonic oscillators, two with a low eigenfrequency (0.7 and 2 Hz) and a high damping ratio (*ζ* > 1), and one with a high eigenfrequency (60 Hz) and a low damping ratio (*ζ* < 1).

### Absence of persistent low-frequency neural oscillations during passive auditory perception

Current theories of speech ^10^ and musical-beat ^30^ perception capitalize on dynamical system theories to describe the interplay between neural and acoustic dynamics. Prior to estimating the capacity of neural oscillations to entrain to sensory stimulations, a clear understanding of the nature of the neural oscillations at play during auditory processing is mandatory. Previous findings have emphasized that rhythmic auditory stimuli principally drive activity at the rate of stimulation, typically in the delta-theta (<8 Hz) range, ^68–70^ and also induce beta-band (∼20 Hz) power modulations ^31, 56, 57^. Thanks to precise stereotactic recordings, we show that rhythmic stimulation induces a complex response at the level of the auditory cortex composed of: (1) evoked activity at the low (∼2.5 and ∼5 Hz harmonic) and high (∼60/80 Hz) rates of acoustic stimulation, combined with transient bursts at tones onset and offset, visible across all frequencies; and (2) induced deactivation in the alpha and beta ranges (7-30 Hz; see Figure 3 and Fig. Supp. 6). Of note, these results, obtained with high-quality sEEG recordings ^53^ and straightforward time-frequency analyses are not subjected to spurious oscillatory artifacts, typically observed after complex signal processing, such as epochs oversampling or phase-amplitude coupling ^71^.

Next, we investigated whether any of the neural responses recorded throughout the cortex present the underdamping property considered as one of the most compelling arguments for entrainment ^45^. Radically, we found that, across frequencies and cortical regions, none of the responses evoked or induced by low-frequency (2.5 Hz) auditory stimulation outlast the stimulus duration. Model-based analyses confirmed that an overdamped oscillator was the best fitting model, *i.e.* an oscillator with no oscillatory behavior after the end of the stimulus, thereby confirming this result.

Different models of neural entrainment describe it either as resulting from the passive mechanistic coupling of neural oscillators with a rhythmic stimulation ^17, 30^, or as an attentional mechanism of sensory selection that involves network-level interactions between higher-order (motor or attentional) regions and sensory cortices ^16, 24, 25, 41, 72^. Our results are thus incompatible with the idea that the auditory cortex can be modelled as an isolated underdamped harmonic low-frequency (delta) oscillator ^30, 43^. They are also inconsistent with the idea that beta-band induced activity entrains at the beat rate during passive listening ^31^.

However, recent studies have shown low-frequency entrainment in auditory regions in task-specific contexts. In particular, in tasks that involve selective attention to one of two presented rhythmic streams ^28, 45^, attention to the moment of occurence of a target ^26^ or attention to the rhythm of a speech stream ^73^. Behavioral studies have also revealed improvements in auditory detection when targets are presented in phase with a rhythmic stimulus, even after several cycles of silence ^30, 74^. Here, we reveal that such a mechanism does not occur during passive perception. To explain the divergence between our results and previous papers that have reported entrainment, we can hypothesize that low-frequency neural entrainment is an active, task-dependent phenomenon. Mechanistic explanations are still lacking, although our results are in line with recent proposals linking entrainment and attention, in which low-order input regions depend on top-down weighting in order to exhibit neural entrainment to input streams ^16, 72, 75^. In such integrative models, top-down systems modulate sensory processing in a proactive and temporally flexible manner to enact entrainment phenomena. Thus, entrainment could reflect selective attention: rhythmic input streams are differentially prioritized, and only the attended stream, being weighted by top-down modulation, entrains oscillations.

A first limitation of this interpretation is that a weak oscillatory response at low-frequency could be masked by a strong omission response elicited at the end of the stimulus. However, we do not see any omission response, probably because we repeated the same auditory stream throughout the experiment, which was hence fully predictable. A second limitation is that low frequencies were carried by the envelope of the stimulus whereas high frequencies were carried by the fundamental frequency of the stimuli. This confounding factor cannot be excluded. However, it is known that amplitude modulated sounds at ∼60/80 Hz produce the same phase-locked evoked responses in the high-gamma range ^76^ that the one we observe. Furthermore, amplitude modulated sounds give rise to a perception of pitch similar to complex tones ^77^. Thus, it is most probable that the frequency range is the key parameter to explain the damping differences we observe between low and high neural oscillations. These results are actually compatible with dynamical system approaches, which model neural population behaviors and provide important insights on the nature of neural oscillations. For example, fundamental differences between high-and low-frequency oscillations have been described. High frequency oscillations are used in the context of small neural ensembles, such as populations of coupled excitatory and inhibitory neurons (PING network), whereas low frequency oscillations usually involve coupled nodes in a network, global ensembles and long-range connections ^78^. Underdamping at such low frequencies is highly unexpected during passive stimulation, whereas it is expected for higher frequency regimes (> 40 Hz).

### Presence of persistent high-frequency neural oscillations throughout the cortex

High frequency phase-locked neural responses to auditory stimulation have been mostly studied at the level of the brainstem ^79, 80^. Recordings of the auditory brainstem responses (ABR) have been developed to assess the integrity of subcortical auditory relays via transient responses to very short sounds (clicks). The use of complex sounds of greater duration has allowed the analysis of a sustained response named frequency-following response (FFR) mimicking the fundamental frequency (and higher harmonics) of the auditory stimulus. Traditionally assessed using a three-electrodes scalp EEG montage, the sources of the ABR, as their name suggests, were considered to be of subcortical origin, notably in the inferior colliculi and medial geniculate bodies ^79, 81–86^. However, the sources of the FFR have recently been subject to intense debate. Two recent papers, using M/EEG ^38^ and fMRI ^39^ have convincingly demonstrated that cortical sources, especially Heschl’s gyrus, also contribute to the scalp-recorded FFR. In this vein, our results confirm the presence of a high-gamma oscillatory response phase-locked to the fundamental frequency of the tones in the auditory cortex. Surprisingly, we also demonstrate that the FFR is actually present in widespread cortical regions, well beyond what was previously observed. Our MEG results are confirmed by the highly spatially-precise and localized sEEG data. The presence of the FFR in such regions could be mediated by the white matter tracts that connect the auditory cortex and different parts of the prefrontal cortex along separate anterior and posterior projection streams ^59^. The fact that a FFR is also present in high-level, integrative cortical regions - such as the motor cortex, supramarginal gyrus, medial temporal lobe or the inferior frontal gyrus-sheds a new light on previous findings showing FFR differences across several types of populations (musical experts, language experts, language impaired populations). For instance, a larger FFR to the fundamental frequency of a sound may well be due to a greater involvement of integrative cortical regions, and may not necessarily imply modifications of subcortical activity via a corticofugal pathway ^87, 88^. Nonetheless, subcortical specificity may be critical with high frequency features such as harmonics or speech formants ^80^.

Crucially, we reveal that the FFR presents an underdamping of up to ten cycles, *i.e.* an oscillatory phase-locked response that persists after stimulus offset. This reflects a passive mechanistic coupling of neural oscillations with a rhythmic stimulation, and is usually modelled with small neural ensembles, such as populations of coupled excitatory and inhibitory neurons (PING network). Thus, the FFR is not only a one-to-one representation of the stimulus, a succession of evoked potentials, but acts as a linear oscillatory filter.

### The damped harmonic oscillator as a model of neural oscillatory activity

The damped harmonic oscillator is standard in physics to study oscillatory phenomena, *e.g.* spring/mass systems, pendulums, torques and electrical circuits. It is defined by a linear second order differential equation, derived from Newton’s second law. Previous works have shown that this model is well-suited to study neural mass dynamics ^67^, *i.e.* spatial averaging of thousands of neurons, in particular in modelling the evoked response ^66^. Although simple and powerful, this model has received little attention in cognitive neuroscience. This lack of interest could arise from the fact that the harmonic oscillator is a phenomenological model, as its parameters capture properties of the neural ensemble and do not refer to physical quantities of the individual neurons, like excitability or conductance ^89, 90^. However, disposing of these biological constraints allows to model with very few parameters the emergent dynamics of the local population, the neural mass, *i.e.* the sEEG signal that we record. In our data, the apparent complexity of the neural response (multiple frequencies, onset and offset responses, harmonics) is in fact reducible to the interaction between the stimulus and a damped harmonic oscillator with three free parameters (*ζ*, 2*πω*_0_ and *Δt*). Furthermore, three clusters of parameters are enough to describe the diversity of cortical responses to a rhythmic auditory stimulation: two overdamped low frequency (0.7 and 2 Hz) and one underdamped high frequency (60 Hz) oscillators. These three clusters show a topology that is consistent with known cerebral networks, namely bilateral auditory cortices, ventral and dorsal auditory pathways ^59^. Given the very limited range of frequencies presented (2.5, 62 and 83 Hz), the interpretation of the 2*πω*_0_ eigenfrequency parameter is limited. The recovered values most probably reflect a mixture of the “true” eigenfrequencies of the recorded neural populations and the stimulus frequencies. The reported values should thus be taken as coarse indexes of “low” versus “high” eigenfrequencies. Refining these values would require presenting a wider range of frequencies.

Concerning the interpretation of the damping parameter, the dichotomy “under-versus overdamping” is usually confounded with the dichotomy “self-sustained oscillator versus superposition of transient event-related potentials”. Indeed, evoked responses are also modelled with oscillators ^91^, and can either stop right after the end of the stimulation (overdamping), due to strong energy dissipation/phase dispersion or continue to oscillate for a while (underdamping). While damping is thus the property that matters from a functional point of view, it can refer to different physiological realities. In particular, two hypotheses yet remain to be clarified: (1) If the neural response is linear, the damping reflects energy dispersion. (2) If the response is non-linear, the damping could also reflect phase dispersion of multiple sustained oscillators. The progressive desynchronization of their phase would induce on average a similar exponential damping.

Finally, it should be noted that the harmonic oscillator is one special case of the broader set of linear filters, widely used in engineering of brain-computer interface. An important objective of this field of research is to define the encoding/decoding function that bridges the stimulus and the brain’s response. Popular models ^65, 92, 93^ are filters, usually approximated by linear regression with regularization due to the large number of fitted parameters. The harmonic oscillator greatly simplifies the regularization problem, as it constrains the space of solution to only three free parameters, without losing explanatory power. Furthermore, analytical solutions are known for any given driving force, which again simplifies the problem by providing solutions with a very low computational cost. Overall, this model is a promising candidate for brain-computer interface engineering, by offering a simple, straightforward encoding/decoding function.

## Methods

### STIMULI AND PARADIGM

#### Stimulus

The auditory stimulus was a bass riff designed to embed both high- (∼60/80 Hz) and low- (2.5 Hz) frequency acoustic modulations, thereby allowing to study properties of both high- and low-frequency neural oscillations (see Figure 1A-C). For a total duration of 6.24 s, it was composed of 16 tones, each lasting 170 ms, presented at a regular pace of 2.56 Hz, with an inter-onset interval of 390 ms. The first eight tones had a fundamental frequency of 83 Hz lasting 14 cycles. The last eight tones had a fundamental frequency of 62 Hz lasting 11 cycles. The two series of eight tones were identical repetition of two complex sounds recorded from an acoustic bass guitar. The acoustic envelope was extracted by computing the absolute value of the Hilbert transform of the signal filtered between 50 and 90 Hz, which captures the fundamental frequencies of both sounds. The inter-stimulus interval was randomly chosen between 2.92, 3.12 and 3.32 s. MEG participants listened to 300 stimuli and sEEG patients 100 stimuli.

#### Stimulus presentation

The stimuli were presented binaurally to participants at an adjusted comfortable level (∼70 dB) using loudspeakers for sEEG patients and Etymotic insert earphones with foam tips (Etymotic Research) for MEG participants. Presentation was controlled with E-prime 1.1 (Psychology Software Tools Inc., Pittsburgh, PA, USA). sEEG patients were passively listening. MEG participants were passively listening and simultaneously watching a silent movie. The whole experiment lasted ∼45 min for MEG participants and ∼15 minutes for sEEG patients.

### COMMON ANALYSES BETWEEN MEG AND sEEG datasets

#### Software

All analyses were done using MNE-python ^94^, FreeSurfer (http://surfer.nmr.mgh.harvard.edu/) and custom scripts written in Python.

#### General strategy

Analysis of sEEG data was done following a fixed effect strategy, *i.e.* considering all patients’ electrodes collectively in an average brain. Individual differences of brain morphology and connectivity were therefore neglected. This strategy was chosen as each patient had a unique electrode implantation map, thus rendering group statistics inappropriate. For MEG data, the strategy was to reduce the normalized signal estimated at the source-level (vertex) to a statistics across participants prior to subsequent analyses, typically a t computed from a one-sample t-test against 0 across participants. Most of the analyses were thereafter consistent between sEEG and MEG data, as having a single average brain with multiple electrodes is similar to having a single average brain with multiple vertices. The only difference was that the sEEG metrics corresponded to z-values, and the MEG metrics to t-values. The fixed-effect strategy we used for the sEEG data has a low generalization power alone. However, replicating the effects in the MEG independent dataset and using a random-effect strategy for the MEG data greatly strengthen our confidence in the generalization of the results.

#### Align-Bin-and-Count (ABC) pipeline: estimation of the duration of a cyclic activity

We developed a method to estimate the number of cycles of oscillatory activity present in a neural signal, relative to the number of cycles present in the stimulus. Note that the same procedure has been used extensively throughout the paper, for high- and low-frequency envelope of the evoked response and inter-trial phase coherence (ITPC). This method has two objectives: find, if any, all responsive sources (sEEG channels or MEG vertices) and estimate the number of activity cycles they exhibit at a specific oscillatory frequency. Persistent activity was defined as >1 oscillatory cycle after stimulus offset. It can be decomposed in 6 steps (see Fig. Supp. 1). The advantage of ABC over more conventional approaches, such as testing ITPC compared to a surrogate distribution, is that it accounts for three potential sources of artifact : 1. the overall power of each source, which can vary greatly (step 1), 2. the lag between stimulus and neural response onsets (steps 2 and 3), and 3. the spurious oscillatory activity (smearing) generated by the filtering process (step 5). Moreover, it is adapted to any kind of signal (evoked response, ITPC or spectral power) and computationally efficient. The six steps are:

1. Z-score the activity of each source across time relative to its prestimulus baseline.
2. Find across all sources the lowest threshold such that none shows onset activity before stimulus onset (data-driven threshold estimation). The threshold is the same for all considered sources, and corresponds to the peak of activity observed during the baseline across all sources.
3. Align each source activity such that time 0 corresponds to the moment where its activity crosses the defined threshold.
4. Discretize the continuous activity in time bins, whose duration depends on the acoustic modulation of interest, *e.g.* 1/83 Hz = 12 ms for the 83 Hz tones; 1/2.5 Hz = 390 ms for the 2.5 Hz stimulus presentation rate.
5. Find across all sources the lowest threshold such that none shows an active bin onset before their onset. We need to define another threshold for the bins because binning implies averaging, rendering the value defined in step (3) irrelevant. Once again, the threshold is the same for all considered sources. This step solves the problem of the smearing induced by the filter: because the filter is symmetric and non-causal, it produces the same artifact before the onset and after the offset. Thresholding to remove spurious activity before the onset also removes spurious activity after the offset.
6. Count the number of consecutive cycles above the threshold defined in step (5). During step (3) and (6), some sources never meet the required criterion and are therefore removed from the analyses. These steps thereby induce a selection of the responsive sources. To validate this procedure, we applied it to the acoustic signal itself and confirmed that it correctly identifies the number of cycles actually present in the stimulus, *e.g.* 11 cycles were estimated for the 62 Hz tones (see Fig. Supp. 2). As the wavelet length decreases with the frequency of interest and that the ABC method uses a binning strategy, all the results are scale free. Therefore, the period duration does not matter for the analysis, as we are counting numbers of cycles and not durations. In this context, there is no specificity of the analysis at 2.6 Hz compared to the analyses at 62 and 83 Hz.

#### Anatomical MRI acquisition and segmentation

The T1-weighted anatomical magnetic resonance imaging (aMRI) was recorded using a 3T Siemens Trio MRI scanner. Cortical reconstruction and volumetric segmentation of participants’ T1-weighted aMRI was performed with FreeSurfer (http://surfer.nmr.mgh.harvard.edu/). This includes: motion correction, average of multiple volumetric T1-weighted images, removal of non-brain tissue, automated Talairach transformation, intensity normalization, tessellation of the gray matter white matter boundary, automated topology correction and surface deformation following intensity gradients. Once cortical models were complete, deformable procedures could be performed including surface inflation and registration to a spherical atlas. These procedures were used to morph current source estimates of each individual for MEG and channel location for sEEG onto the FreeSurfer average brain for group analysis.

### STEREOTACTIC EEG (sEEG)

#### Participants

16 patients (5 females, mean age 26.9 y, range 9-46 y, see Table Supp. 1) with pharmacoresistant epilepsy took part in the study. They were implanted with depth electrodes for clinical purposes at the Hôpital de La Timone (Marseille, France). Neuropsychological assessments carried out before sEEG recordings indicated that all participants had intact language functions and met the criteria for normal hearing. None of them had their epileptogenic zone including the auditory areas as identified by experienced epileptologists. Patients provided informed consent prior to the experimental session, and the study was approved by the Institutional Review board of the French Institute of Health (IRB00003888).

#### Data acquisition

Depth electrodes (0.8 mm, Alcis, Besançon, France) containing 10 to 15 contacts were used to perform the functional stereotactic exploration. Contacts were 2 mm long and spaced from each other by 1.5 mm. The locations of the electrode implantations were determined solely on clinical grounds. During the recording session, participants laid comfortably in a chair in a sound attenuated room. sEEG signals were recorded at a sampling rate of 1000 Hz using a 256-channels BrainAmp amplifier system (Brain Products GmbH, Munich, Germany) and bandpass filtered between 0.3 and 500 Hz. A scalp electrode placed in Fz was used as the recording reference.

#### Anatomical localization of electrodes

Anatomical localization of electrodes was performed using a local software developed at the Hôpital de La Timone ^95^. First, an automatic procedure coregistrate electrodes location on the scanner and patient’s MRI. Second, the morphing matrix computed by Freesurfer used to project one individual’s brain onto the average brain was applied directly on these locations.

#### Pre-processing

In order to remove power line artifacts, we first applied a notch filter at 50 Hz and harmonics up to 300 Hz. The signal was then re-reference using a bipolar montage, *i.e.* activity of each channel was subtracted from its following neighbour on the electrode. The continuous signal was then epoched from -1 to 9 s relative to the onset of each stimulus. Such long epochs allows to study both the response during the stimulus, from 0 to 6.24 s, and the silence directly following stimulus offset, from 6.24 up to 9 s. No baseline correction was applied, as effects potentially present in the silence can leak to the baseline of the next trial and therefore be removed by any process of baseline correction. For each source, epochs with ±500 μV peak-to-peak amplitude artifacts were rejected. Sources with > 70% rejected epochs were entirely removed, as they most likely contained epileptic discharges. Finally, sources showing low voltage epileptic activity were removed based on visual inspection. Overall, 2648/3139 sources were kept, with on average 82/100 epochs (see Table Supp. 1).

#### Time-frequency decomposition

Trial-by-trial time-frequency decomposition was carried out in a range of 100 frequencies, logarithmically ranging from 2 to 150 Hz. Morlet wavelet transform was applied to the data using the MNE-python function *time_frequency.tfr_morlet*, with parameter n_cycles = 6. From the resulting complex representation, both inter-trial phase coherence (ITPC) and power were extracted as follows: 

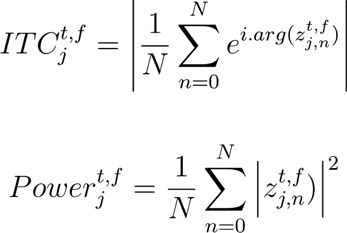

Where 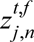 designates the complex time-frequency representation of source j at trial n over N, frequency f and time t. Similarly, 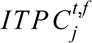 designates the ITPC of source j at frequency f and time t, and 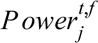 its power.

#### Frequency following response analysis

In order to study the FFR, we re-epoched the signal from -0.05 to 0.35 s relative to the onset of each tone. We computed the evoked activity as the mean across epochs. In order to study only the high-frequency component of the evoked response, we filtered it between 82 and 84 Hz for the 83 Hz tones and between 61 and 63 Hz for the 62 Hz tones, using a one-pass, zero-phase, non-causal bandpass FIR filter with a hamming window (see Fig. Supp. 1 and Fig. Supp. 2 for the impulse response of the filter). We then extracted the envelope as the absolute value of the Hilbert transform, and applied our ABC pipeline (see **COMMON ANALYSES**) to select the responsive sources and estimate the duration of their activity relative to the number of cycles present in the stimuli (14 for the 83 Hz tones and 11 for the 62 Hz tones). The automatically selected thresholds for the onsets (step 3) were 5.0 z-score for the 83 Hz tones and 4.9 for the 62 Hz tones. The automatically selected thresholds for the bins activity (step 5) were 5.1 for the 83 Hz tones and 4.8 for the 62 Hz.

#### Envelope-following response analysis

We apply the same line of reasoning to investigate the EFR at 2.5 Hz. We computed the evoked activity as the mean across all epochs. We filtered this activity between 1.6 and 3.6 Hz and extracted the envelope. We then apply ABC on this envelope signal (see **COMMON ANALYSES**). The selected thresholds were 5.4 for the onsets (step 3) and 4.3 for the bins (step 5).

#### Definition of canonical frequency bands based on auditory cortex activity

Localization of the auditory cortex was based on functional criteria, including latencies and shape of the auditory evoked potentials to pure tones, tested on an independent session. Primary auditory cortex (PAC) was defined by the presence of a P20/N30 complex, and secondary auditory cortex (SAC) by the presence of a P40/N50 complex ^96, 97^. Responses of PAC and SAC were averaged, as no main differences were found in any of our analyses. Definition of canonical frequency bands was then based on the profile of the spectrum of ITPC and power (averaged across time during the stimulus duration, from 0 to 6.24 s). It resulted in an empirical definition of delta (2-3.5 Hz), theta (4-7 Hz), alpha (8-11 Hz), and gamma bands (50-110 Hz). Although not prominent neither in the sEEG dataset nor in the MEG dataset, we added a beta band (12-22 Hz) based on existing literature. Indeed, previous works have shown coupling of the beta band power with the phase of the stimulus envelope at similar paces ^31, 55^.

#### Inter-trial phase coherence fluctuations at 2.5 Hz

Inter-trial phase coherence (ITPC) was first averaged across frequencies inside each frequency band (delta, theta, alpha, beta, and gamma). For each band we then applied the ABC pipeline (see **COMMON ANALYSES**) to estimate responsive sources as well as the duration of significant oscillatory activity. Thresholds for onsets (step 3) were 3.4 (delta), 4.2 (theta), 5.4 (alpha), 6.3 (beta), 8.2 (gamma). Thresholds for the bins (step 5) were respectively 3.2, 3.7, 3.8, 3.4, 3.6.

#### Temporal response function analysis

To analyse power signals, we relied on a recently developed powerful methodology that estimates temporal response functions (TRF) by relying on encoding/decoding models of electrophysiological activity ^65^. TRF rely on the assumption that the activity can be expressed as a linear convolution between the input stimulus and a filter. The filter is typically unknown and therefore estimated by a least-square ridge regression. We reasoned as follows: if the power is modulated by the envelope of the auditory stimulus, then these fluctuations are informative about the stimulus dynamics. We should therefore be able to train a TRF to decode the envelope based on the power information. If the power continues to exhibit these fluctuations in the silence directly following the offset of the stimulus, we should still be able to decode the envelope oscillations during this post-stimulus period. Furthermore, and contrary to the ABC method, the TRF would reveal effects that are suppressed by baseline correction, as it does not involve baseline correction. We trained a TRF on all available power information, namely power in all epochs, sources, frequencies and time points, to decode the envelope of the auditory stimulus. We trained the model on the second part of the auditory stimulus, from 3.12 to 5.85 s after stimulus onset (which corresponds to the presentation of the six 62 Hz tones), and evaluated it on the first part of the stimulus from 0.39 to 2.73 s (which corresponds to the presentation of the six 83 Hz tones), and on the silence directly following, from 6.24 to 8.58 s. The time windows were chosen to have the same length, and to avoid the strong evoked activity present for the first tone and the changing tone. Performance was defined as the coefficient of determination (R^2^) between the predicted envelope and the actual envelope. The ridge parameter (*α* = 1000) was chosen to maximize the performance of the TRF on the signal during the first part of the stimulus. In order to reduce dimensionality and improve computational speed, we fitted a principal components analysis (PCA) prior to the TRF training. To avoid confounds due to this PCA procedure, we fitted it on the power in the second part of the stimulus and applied it without any further adjustments on the power in the first part and in the silence. We kept 60 components, explaining 99% of the total variance.

#### Damped harmonic oscillator model

We fitted the sEEG data to a damped harmonic oscillator model. The model is described by the following linear differential equation:

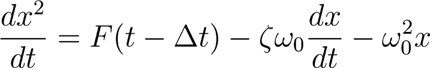

We used a grid search approach to find optimal parameters. For each set of parameters, we simulated 100 epochs of the harmonic oscillator time course with random initial conditions, with F being our auditory stimulus. Although analytically solved for any function F, we used a numerical solution (Euler’s method with a time step of 1/40000 s) because our stimulus has no simple functional form. In order to compare model time courses and sEEG data, we apply the same processing steps: band-pass filter, resampling at 1000 Hz, Morlet wavelet transform, extraction of power and ITPC. Goodness-of-fit was then measured as the coefficient of determination (R^2^) of the linear regression between the model ITPC (or power) and the sEEG data ITPC (or power). The grid search explored a wide range of values: *ζ* logarithmically spaced values between 0.01 and 100 in 25 steps, 2*πω*_0_ logarithmically spaced values between 0.1 and 100 Hz in 25 steps, *Δt* linearly spaced values between 0 and 400 ms in 20 steps. This procedure was used to fit all sEEG channels and output a parameters matrix of size n_channels x 3. This matrix was used to cluster the electrodes using the k-means algorithm. The optimal number of clusters (k=3) was selected using the silhouette index, a standard measure that compares the mean intra-cluster distance and the mean nearest-cluster distance for each sample. Contrary to the ABC method, this modelling approach would reveal effects that are suppressed by baseline correction, as it does not involve baseline correction.

### MAGNETOENCEPHALOGRAPHY (MEG)

#### Participants

We collected data from 15 participants (8 females, median age 27 y, age range 23 to 40 y) after providing informed consent. All had normal hearing, reported no neurological deficits and received 40 euros for their time. The experiment was approved by the National Ethics Committee on research on human subjects.

#### Data acquisition

MEG data were recorded with a whole head 4D-neuroimaging system with 248 magnetometers. Participants were laying in horizontal position under the MEG dewar, facing a screen displaying a silent movie. MEG recordings were sampled at 678 Hz and bandpass filtered between 0.3 and 500 Hz. Four head position coils (HPI) measured the head position of participants before each block. Prior to the session, 2 minutes of empty room recordings were acquired for the computation of the noise covariance matrix.

#### Pre-processing

In order to remove power line artifacts, we first applied a notch filter at 50 Hz and harmonics up to 300 Hz. We further low-pass filter the signal below 150 Hz and resampled it at 500 Hz. An independent components analysis (ICA) was performed on the band-pass signal between 1 and 30 Hz and components exhibiting topographical and time courses signatures of eye blinks or cardiac artifacts were removed from the data. The continuous signal was then epoched from -1 to 9 s relative to the onset of each stimulus. No baseline correction was applied. Epochs with ±5 pT peak-to-peak amplitude artifacts were rejected (on average 4.6 %).

#### MRI-MEG co-registration and source reconstruction

The coregistration of MEG data with the individual’s structural MRI was carried out by realigning the digitized fiducial points with MRI slices. Using MRILAB (Neuromag-Elekta LTD, Helsinki), fiducials were aligned manually on the MRI slice. Individual forward solutions for all source reconstructions located on the cortical sheet were next computed using a 3-layers boundary element model ^98, 99^ constrained by the individual aMRI. Cortical surfaces were extracted with FreeSurfer and decimated to about 10,240 sources per hemisphere with 4.9 mm spacing. The forward solution, noise and source covariance matrices were used to calculate the depth-weighted (default parameter *γ* = 0.8) and noise-normalized dynamic statistical parametric mapping (dSPM) ^100^ inverse operator. This unitless inverse operator was applied using a loose orientation constraint on individuals’ brain data by setting the transverse component of the source covariance matrix to 0.2 (default value). The reconstructed current orientations were pooled by taking the norm, resulting in manipulating only positive values. The reconstructed dSPM estimates time series and time-frequency plane were morphed onto the FreeSurfer average brain for group analysis and common referencing.

#### Time-frequency decomposition

Trial-by-trial time-frequency decomposition was performed using the same procedure and same parameters as for the sEEG.

#### Frequency following response analysis

As for sEEG, we epoched data, computed the evoked response, filtered it at the stimulus fundamental frequency, extracted the envelope and applied a z-score normalisation. We applied a t-test against 0 across participants at each time points and each sources. We then applied the ABC pipeline (see **COMMON ANALYSES**) to select the responsive sources and estimate the duration of their activity. Thresholds for MEG were manually fixed, and corresponded to p < 0.01 (t-statistic ∼2.7, df = 14).

#### Low frequency amplitude analysis

As for sEEG, we computed the evoked response, filtered it at the stimulus fundamental frequency, extracted the envelope and applied a z-score normalisation. Then, we applied a t-test against 0 across participants at each time points and each sources. We then followed the ABC pipeline (see **COMMON ANALYSES**) with a p < 0.01 threshold.

#### Anatomical localization of auditory cortex

We relied on an anatomical criterion to localize auditory cortex. We used the Destrieux atlas (Destrieux et al., 2010), label *temporal transverse*, comprising primary and secondary auditory cortices (left hemisphere “S_temporal_transverse-lh”: 564 vertices, right hemisphere “S_temporal_transverse-rh” 413 vertices). We first applied a singular value decomposition (SVD) to the time courses of each source within the label and used the scaled and sign-flipped first right-singular vector as the label time course. The scaling was performed such that the power of the label time course was the same as the average per-vertex time course power within the label.

#### Inter-trial phase coherence fluctuations at 2.5 Hz

Similarly to sEEG, ITPC was first averaged across frequencies inside each frequency band, for each participant, source, frequency band and time point. We z-score the signal across time compared to its baseline. We then applied a t-test against 0 across participants. We used the ABC pipeline (see **COMMON ANALYSES**) as previously, with a p < 0.01 threshold.

#### Temporal response function analysis

The TRF model was fitted on sensor space in MEG to reduce computational load. Furthermore, sensors contain the same amount of information as sources, thus estimating TRF on sensors or on sources is theoretically equivalent. We therefore trained the model on all available power information, namely power in all participants, sensors, epochs, frequencies and time points, to decode the envelope of the auditory stimulus. The ridge parameter (*α* = 1.0x10^8^) was chosen to maximize the performance of the TRF on the signal during the first part of the stimulus (as for sEEG). In order to reduce dimensionality and improve computational speed, we fitted a principal components analysis (PCA) prior to the TRF training. We kept 101 components, explaining 99% of the total variance.

## Corresponding Author and Lead Contact

Jacques Pesnot Lerousseau, Aix-Marseille Univ, INS, Inst Neurosci Syst, Marseille, France;

## Acknowledgments

We thank C. Barbosa and P. Marquis for helping with the data acquisition, R. Zatorre for insightful comments, S. Baillet, D. Battaglia, C. Bernard, V. Jirsa and all colleagues from the Institut de Neuroscience des Systèmes for useful discussions.

## Author contributions

Conceptualization D.S; Data curation J.P.L.; Formal Analysis J.P.L.; Funding acquisition D.S.; Investigation J.P.L.; Methodology J.P.L., B.M. and D.S.; Project administration D.S.; Resources A.T. and D.S.; Software D.S.; Supervision B.M. and D.S.; Validation A.T., B.M. and D.S.; Visualization J.P.L.; Writing – original draft J.P.L., B.M. and D.S.; Writing – review & editing J.P.L., A.T., B.M. and D.S.

## Supplementary Figures

**Supplementary Figure 1.**
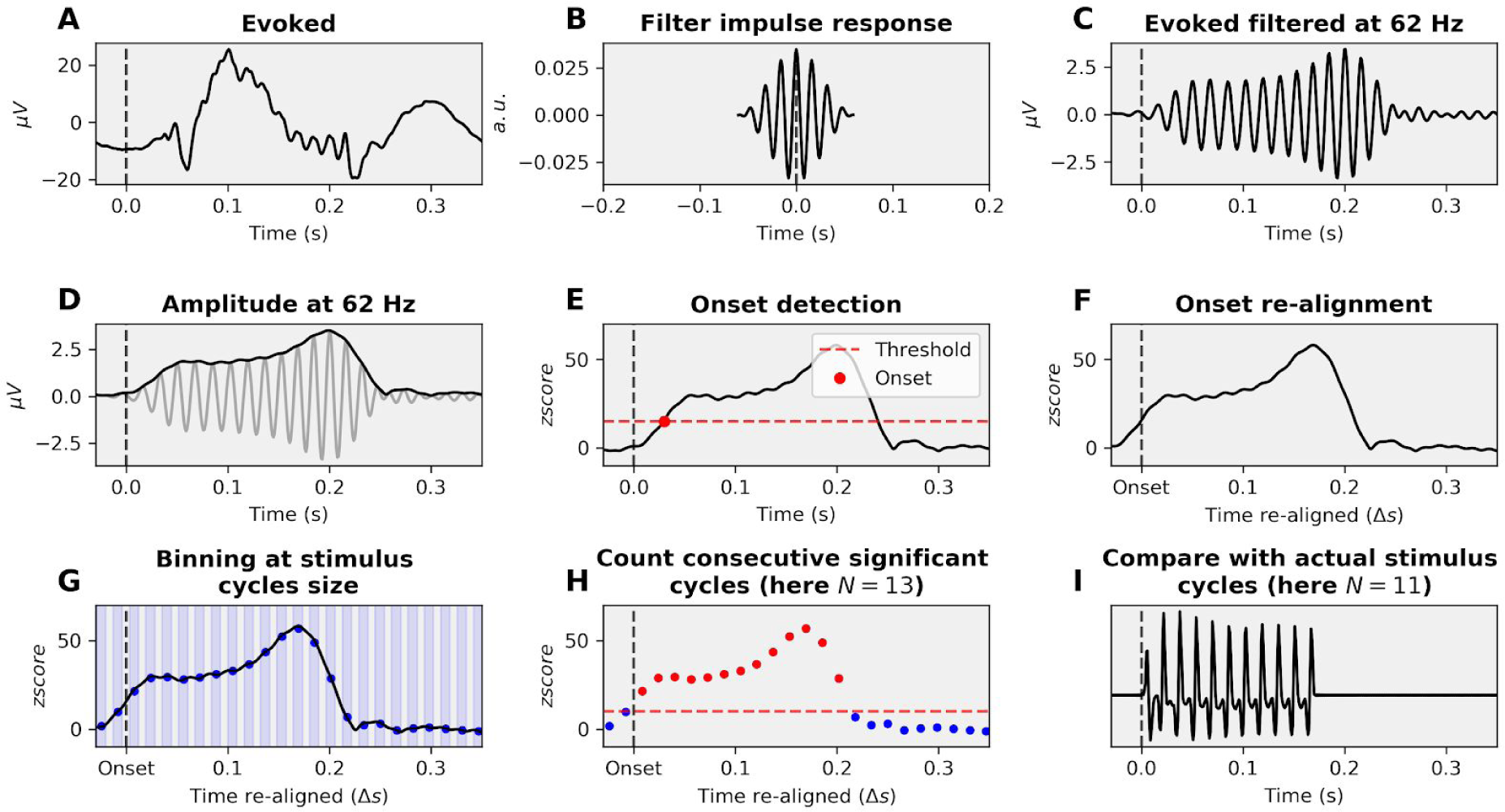
Binning methodology. **A.** Signal of interest, here an evoked response. **B.** Impulse response of the one-pass, zero-phase, non-causal bandpass FIR filter with a hamming window. **C.** Filtered signal at the frequency of interest, here 62 Hz. **D.** Extraction of the envelope of the signal, as the absolute value of the Hilbert transform. **E.** Onset detection. The threshold is chosen such that no channel/vertice shows activity before stimulus onset. **F.** Signal re-alignment. **G.** Binning of activity at the relevant window size (here 1/62 = 16 ms). **H.** Consecutive significant cycles count. Another threshold is chosen such that no bin of any channel/vertice is significant before onset. **I.** Comparison between the stimulus number of cycles and the signal number of cycles. Here, the evoked activity shows 2 cycles of activity more than the stimulus.

**Supplementary Figure 2.**
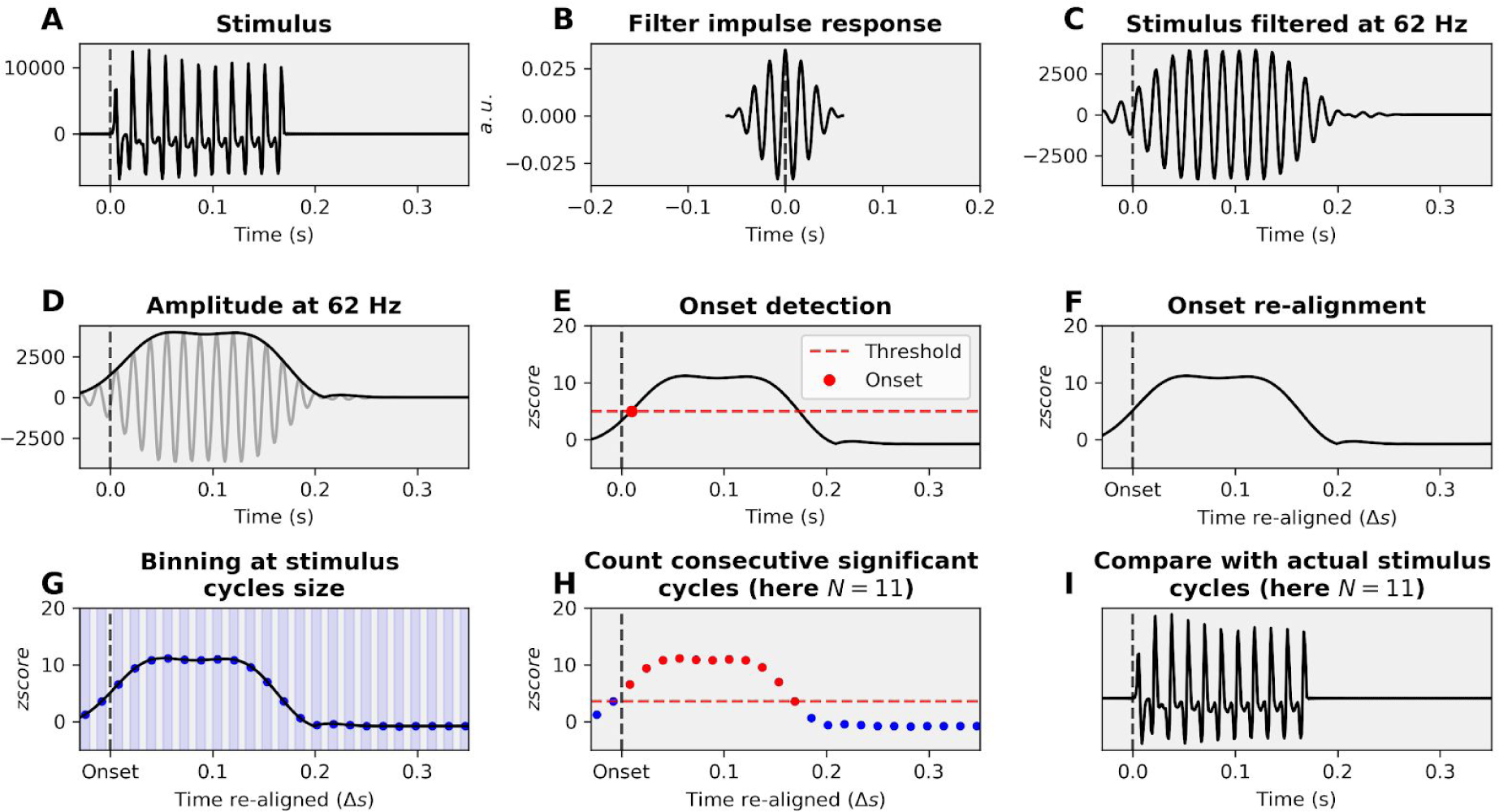
Validation of the binning methodology. **A.** Control signal of interest, here the stimulus itself. **B.** Impulse response of the one-pass, zero-phase, non-causal bandpass FIR filter with a hamming window. **C.** Filtered signal at the frequency of interest, here 62 Hz. **D.** Extraction of the envelope of the signal, as the absolute value of the Hilbert transform. **E.** Onset detection. The threshold is chosen such that no channel/vertice shows activity before stimulus onset. **F.** Signal re-alignment. **G.** Binning of activity at the relevant window size (here 1/62 = 16 ms). **H.** Consecutive significant cycles count. Another threshold is chosen such that no bin of any channel/vertice is significant before onset. **I.** Comparison between the stimulus number of cycles and the control signal number of cycles. Here, the method correctly identifies 11 cycles in the 11 cycles stimulus.

**Supplementary Figure 3.**
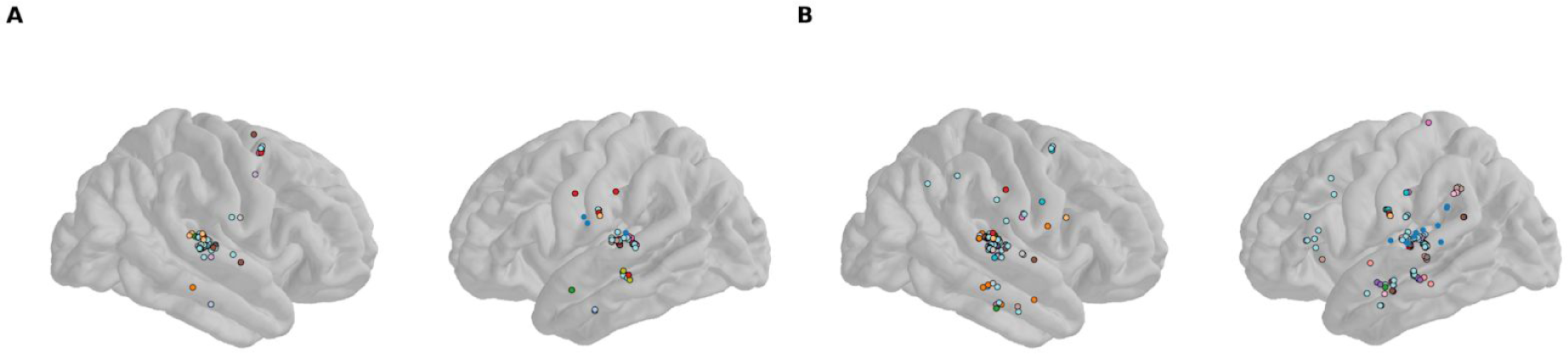
Individual localisation of the sEEG channels showing a FFR during the stimulus. **A.** Individual localization of sEEG channels showing a FFR for the 83 Hz tones. Only channels with activity outlasting stimulus duration are shown. Color represents participants (N = 16). **B.** Same for the 62 tones. Colors representing participants are the same as in panel A.

**Supplementary Figure 4.**
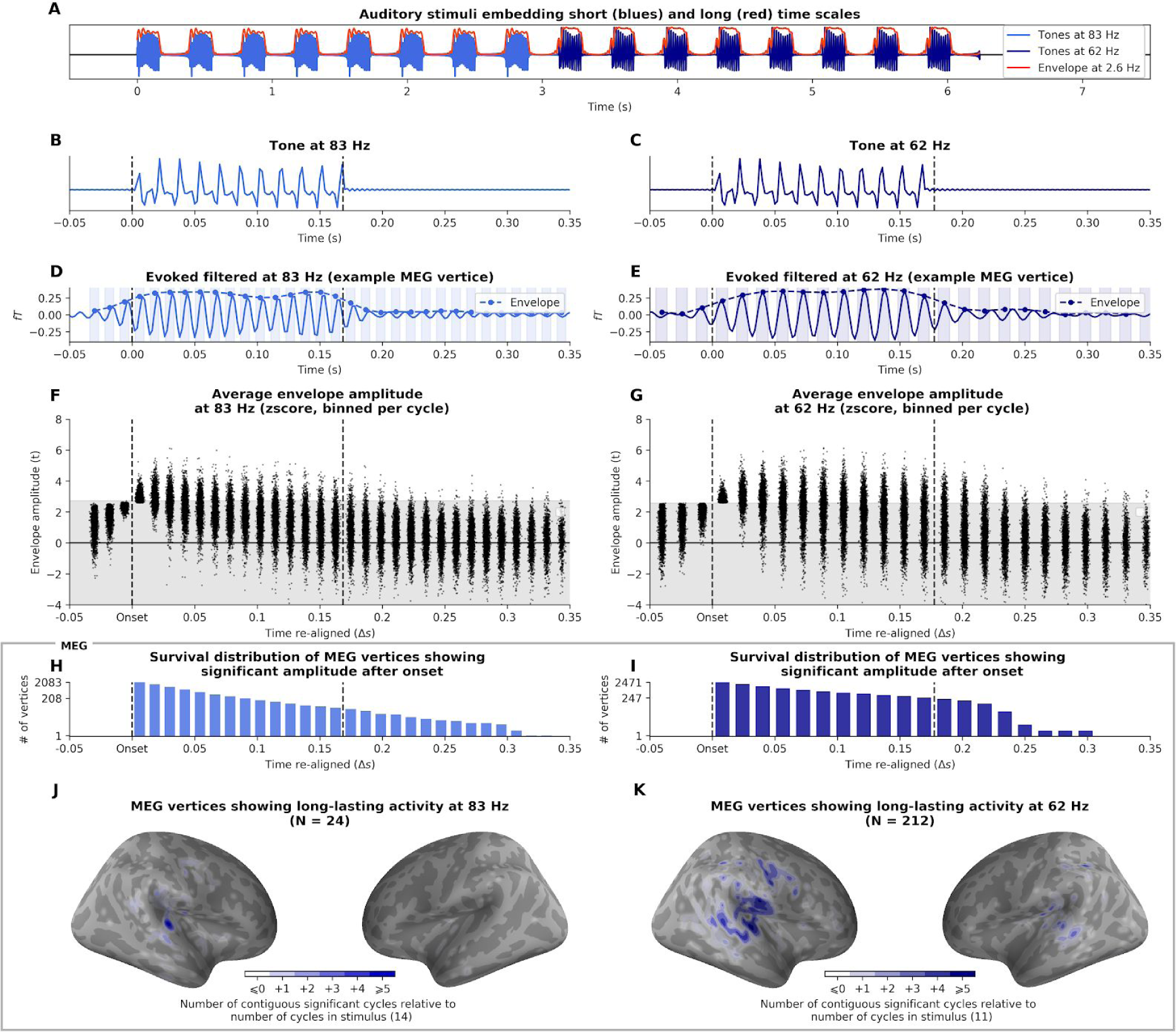
Estimation of frequency-following response duration (FFR, 83 and 62 Hz) in MEG. **A.** Auditory stimulus of 8 tones with high frequency carriers at 83 Hz followed by 8 tones at 62 Hz presented at a 2.5 Hz rate (envelope). **B-C.** Waveform of the 83 and 62 Hz tones, made-up of 14 and 11 oscillatory cycles respectively. **D-E.** Filtered evoked response at 83 and 62 Hz from an example MEG vertice. Dotted line represents the envelope of the neural signal. Shaded areas illustrates the binning window of the ABC pipeline (see Methods). **F-G.** Distribution of binned amplitude of the filtered evoked response across all MEG vertices that have a FFR. Time 0 indicates onset response, as signals are re-aligned prior to the binning process. **H-I.** Survival distribution of MEG vertices with consecutive significant activity after onset. 24 and 212 vertices outlast the 0.170 s duration of the 83 and 62 Hz stimuli, respectively. **J-K.** Localization of MEG vertices showing a FFR. Color intensity indicates the number of cycles outlasting stimulus duration.

**Supplementary Figure 5.**
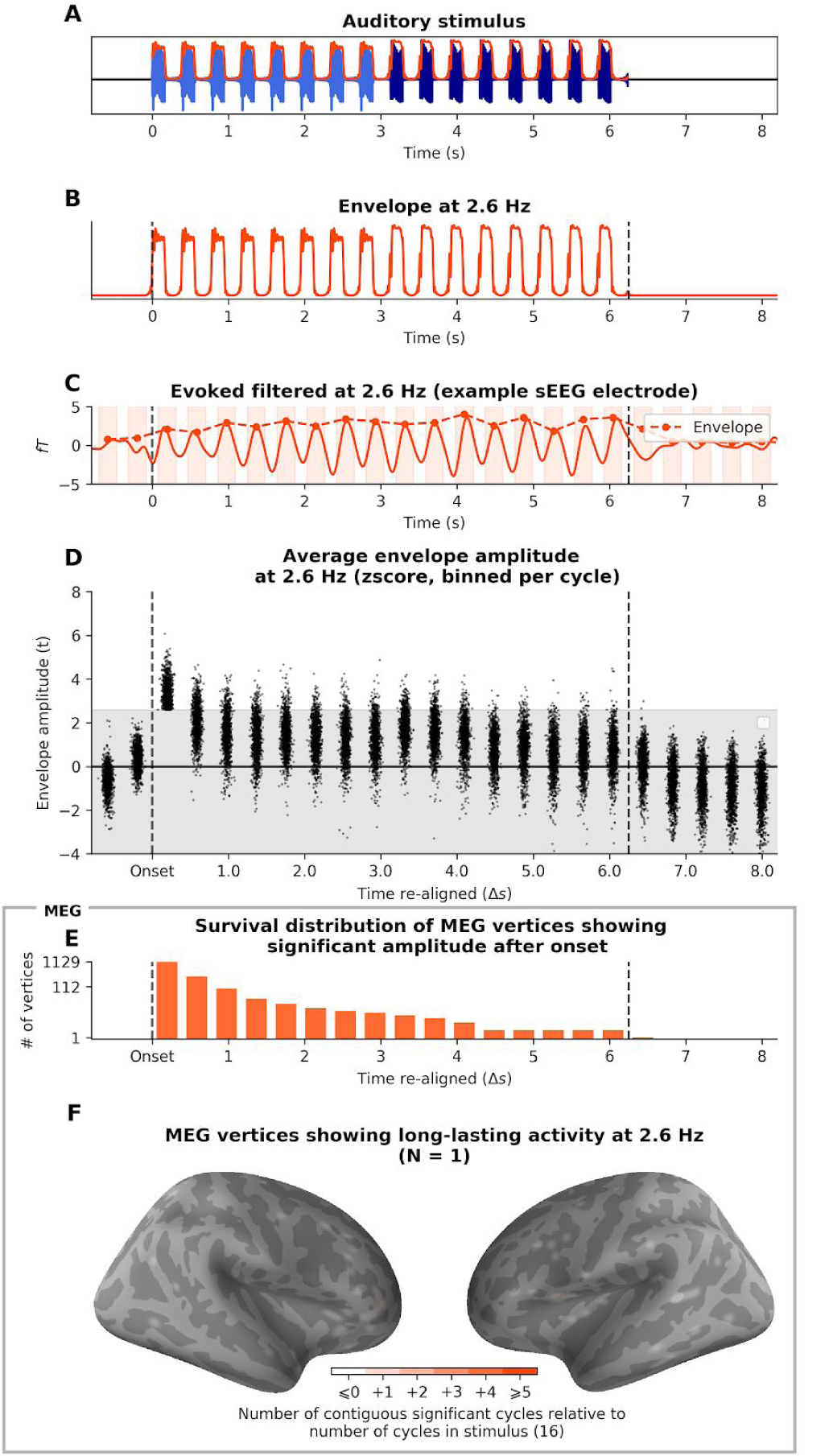
Estimation of envelope-following response duration (EFR, 2.5 Hz) in MEG. **A.** Auditory stimulus. **B.** Waveform of the envelope fluctuation at 2.5 Hz. **C.** Filtered evoked response at 2.5 Hz of an example MEG vertice. Dotted line represents the envelope of the neural signal. Shaded areas illustrates the binning process of the ABC pipeline (see Methods). **D.** Distribution of binned amplitude of the filtered evoked response across all MEG vertices that have an EFR. Time 0 indicates onset response, as signals are re-aligned prior to the binning process. **E.** Survival distribution of MEG vertices activity showing consecutive significant activity after onset. Vertices whose activity lasts more than 16 cycles outlast the duration of the stimulus (N = 1). **F.** Localization of MEG vertices that have an EFR. Color intensity indicates the number of cycles of post-stimulus activity.

**Supplementary Figure 6.**
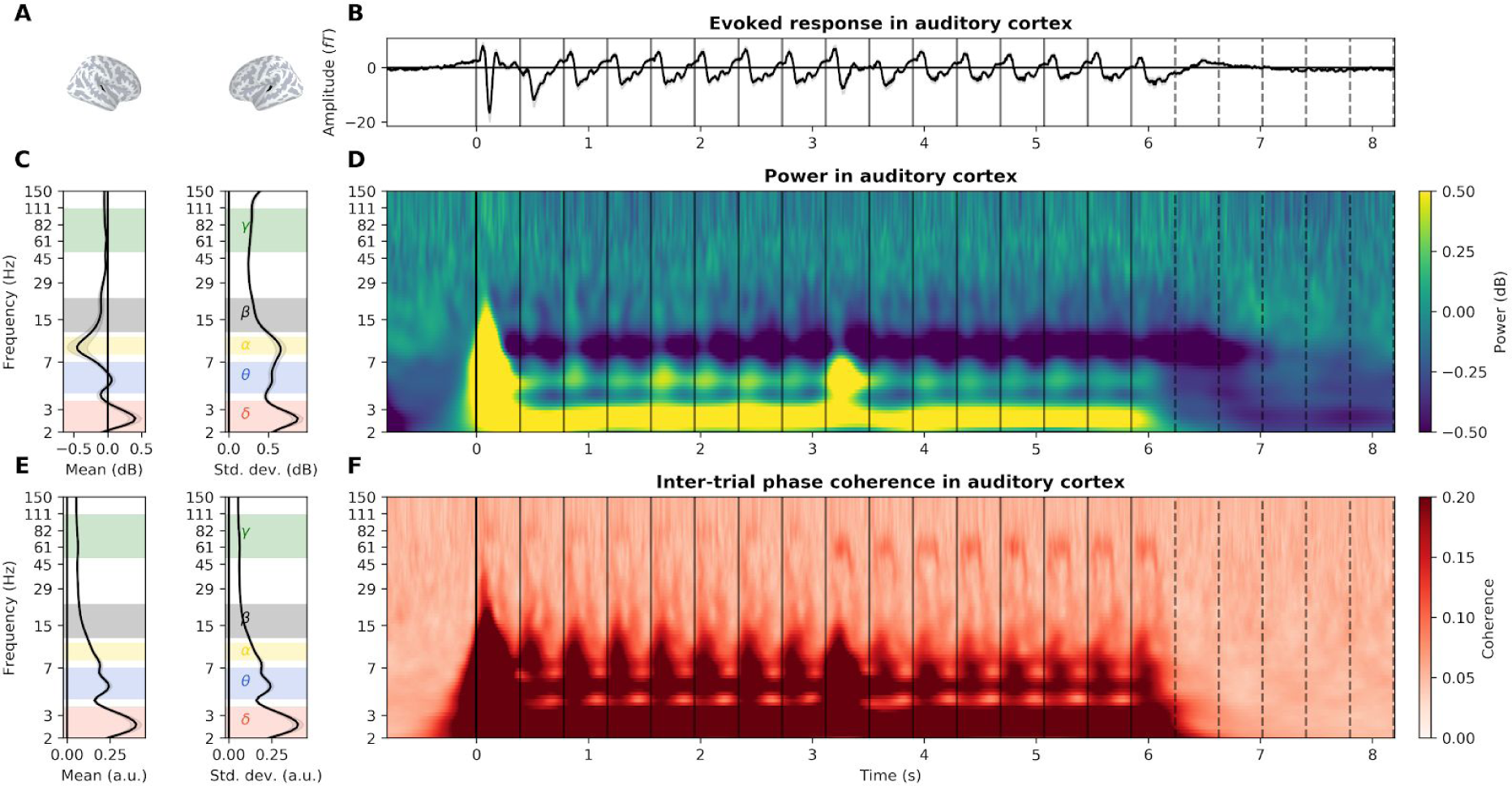
Detailed envelope-following response in the auditory cortex recorded with MEG. **A.** Localization of the MEG vertices located in bilateral auditory cortices. **B.** Evoked response. Grey shaded area represents the standard error of the mean (s.e.m.) across participants. Vertical plain and dotted lines respectively indicate the onset of each tone and their theoretical continuation in the silence. **C**. (Left) Average and (right) standard deviation of power over time during stimulus presentation (0 - 6.2 s). Selected frequency bands are indicated by colored shaded areas (δ delta: 2-3.5 Hz, θ theta: 4-7 Hz, α alpha: 8-11 Hz, β beta: 12-22 Hz, γ gamma: 50-110 Hz). **D.** Power, averaged across participants, in dB relative to baseline. **E.** (Left) Average and (right) standard deviation of inter-trial phase coherence (ITPC) over time during the presentation of the stimulus. **F.** Inter-trial phase coherence (ITPC), averaged across participants.

**Supplementary Figure 7.**
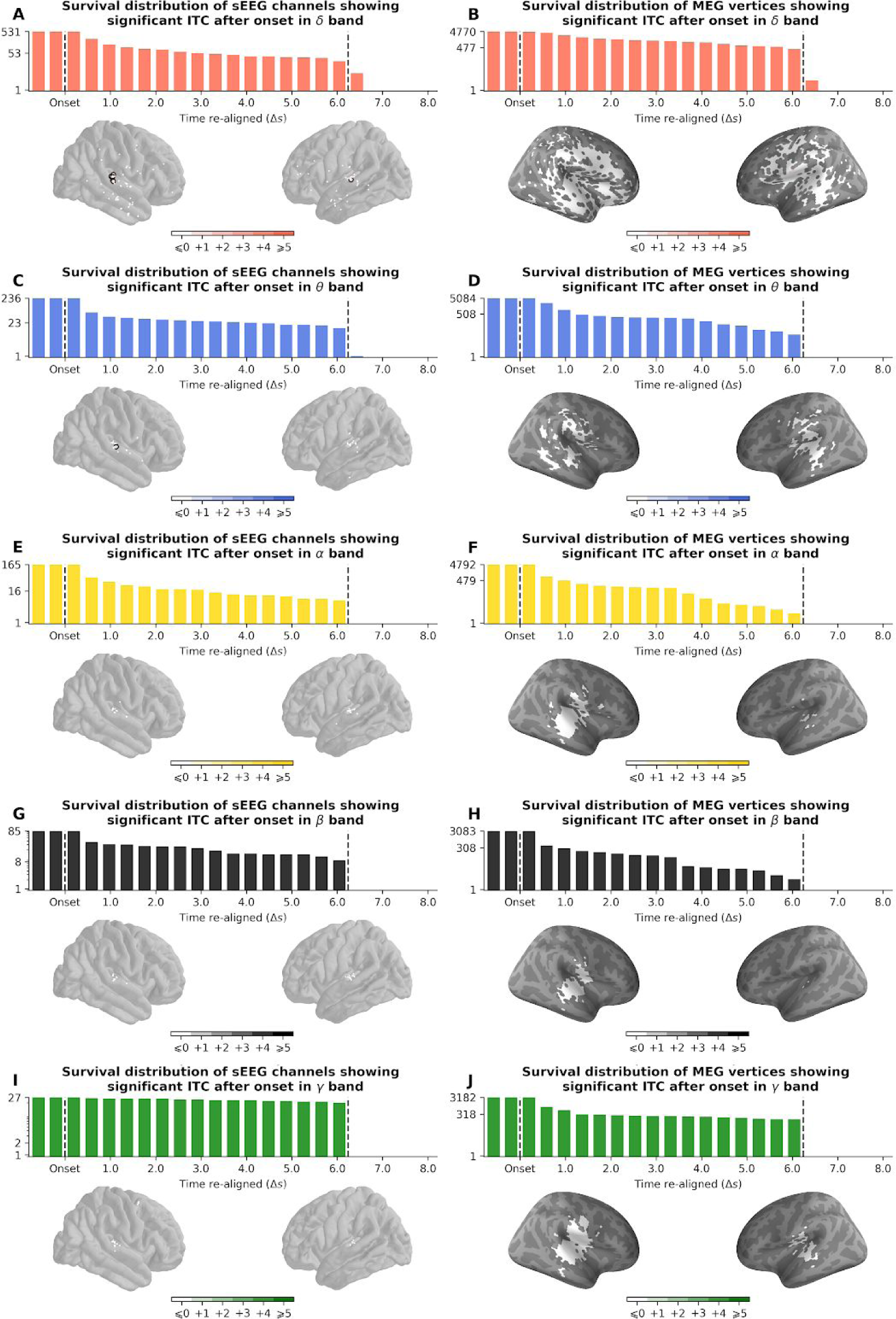
Estimation of duration of inter-trial phase coherence (ITPC) duration per frequency bands. **A.** (Top) Survival distribution of sEEG channels activity showing consecutive significant ITPC in delta band after onset. Channels still showing ITPC after 16 cycles show entrainment (N = 6). (Bottom) Localization of responsive sEEG channels. Color intensity indicates the number of cycles their activity exceed those of the stimulus. **B.** (Top) Survival distribution of MEG vertices activity showing consecutive significant ITPC in delta band after onset. (Bottom) Localization of responsive MEG vertices. Same color code as in A. **C-D.** Same for theta band. **E-F.** Same for alpha band. **G-H.** Same for beta band. **I-J.** Same for gamma band.

**Supplementary Figure 8.**
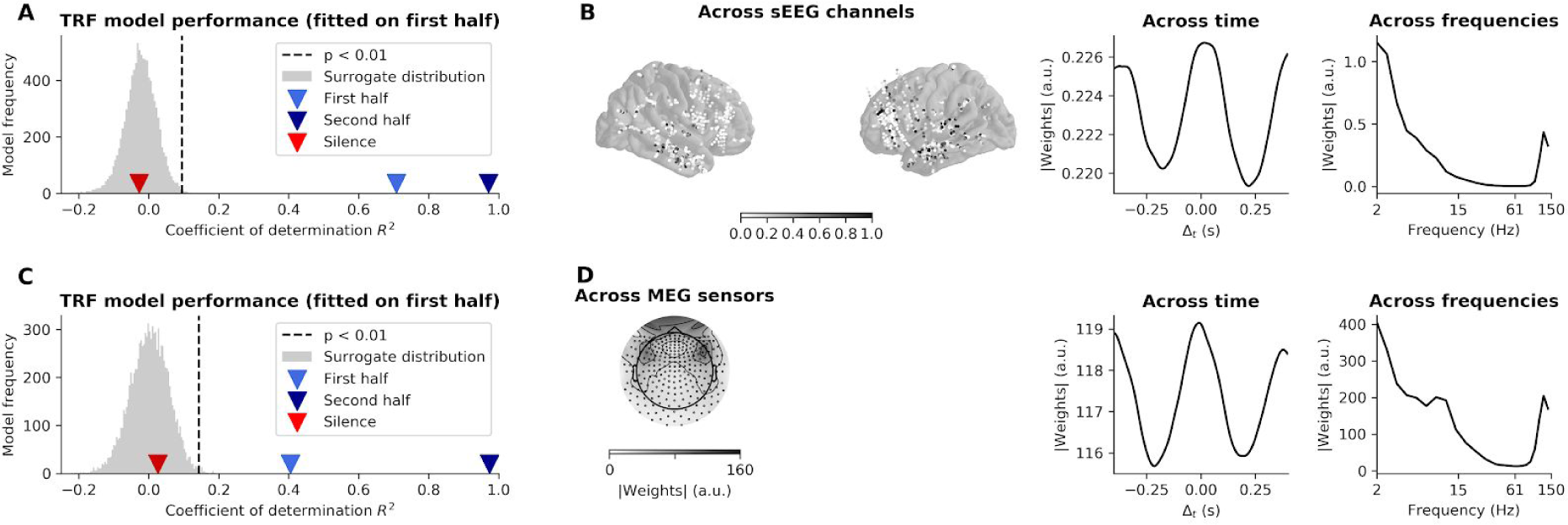
Decoding of the stimulus envelope in sEEG and MEG power data. **A.** Performance of the decoding model (Temporal Response Function, TRF), trained on the power signal (2-150 Hz) of sEEG data recorded during presentation of the second half of the stimulus (62 Hz tones) and generalized to the first half of the stimulus (83 Hz) tones and the post-stimulus silence. Performance is assessed by the coefficient of determination R^2^ obtained between the envelope reconstructed from the neural data and the actual stimulus envelope. Grey distribution indicates performance of TRF model on surrogate data. **B.** Marginal distributions of TRF weights (in absolute value) of sEEG data. Values are averaged in the sEEG channels (left), time (middle) and frequencies (right). **C.** Performance of the decoding model trained on MEG data. **D.** Marginal distributions TRF weights (in absolute value) of MEG data.

**Supplementary Figure 9.**
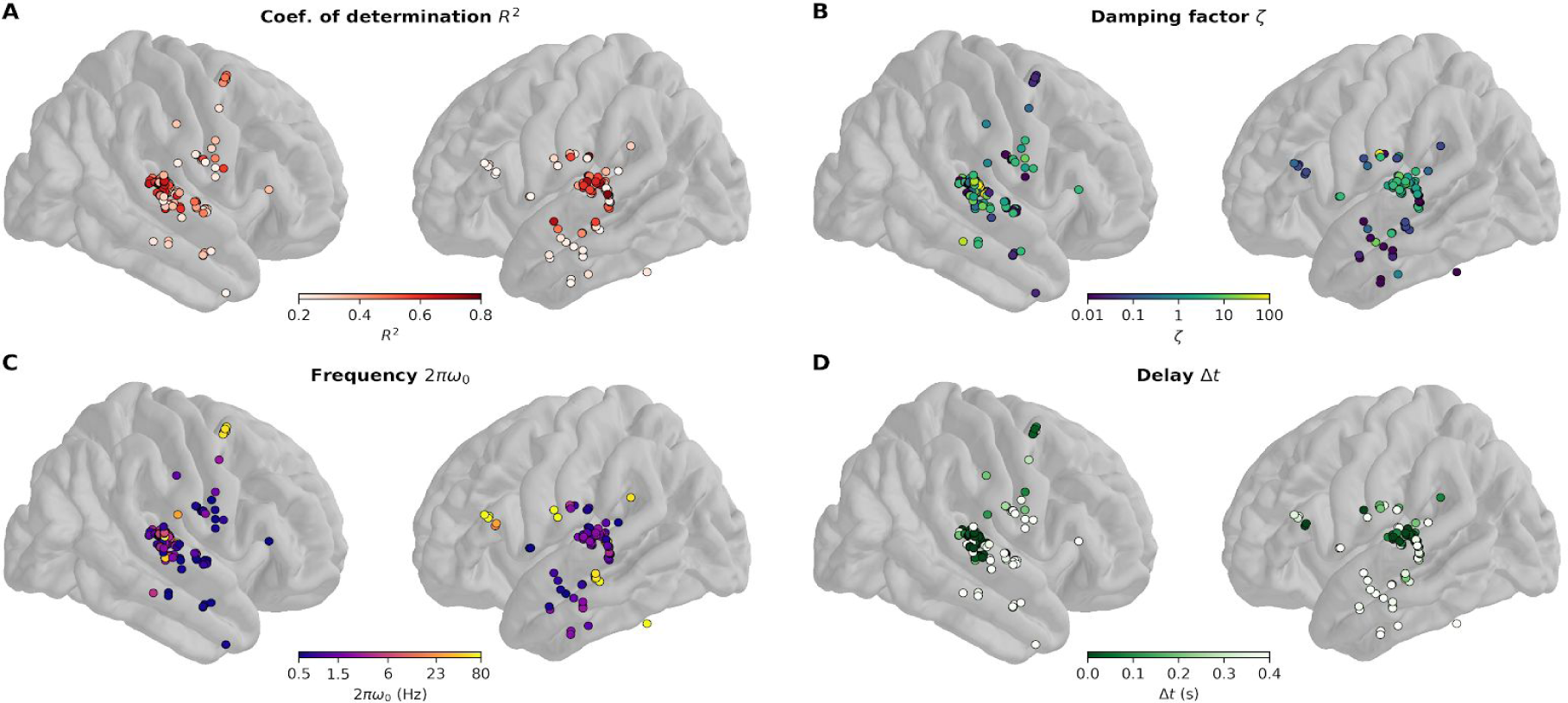
Damped harmonic oscillator fit to each sEEG channels. **A.** Coefficient of determination of the linear regression between ITPC of the best fitting model and ITPC of the data. **B.** Damping ratio *ζ* of the best fitting model. **C.** Eigenfrequency 2*πω*_0_ of the best fitting model. **D.** Time delay *Δt* of the best fitting model. Only channels with R^2^ > 5% are shown.

**Supplementary Table 1.** sEEG participants characteristics. For each participants (N = 16), the table displays the total number of channels (in total 3139), the number of channels kept in the analyses (in total 2648), the average number of epochs kept in the analyses (grand average 82 %), the number of selected channels in the right primary and secondary auditory cortices (in total 13, see Methods) and the number of selected channels in the left primary and secondary auditory cortices (in total 12, see Methods).

| Participant | 1 | 2 | 3 | 4 | 5 | 6 | 7 | 8 | 9 | 10 | 11 | 12 | 13 | 14 | 15 | 16 |
| --- | --- | --- | --- | --- | --- | --- | --- | --- | --- | --- | --- | --- | --- | --- | --- | --- |
| Total channels | 189 | 224 | 189 | 249 | 206 | 150 | 189 | 224 | 189 | 214 | 184 | 160 | 209 | 192 | 180 | 191 |
| Analyzed channels | 185 | 218 | 183 | 219 | 198 | 139 | 188 | 102 | 188 | 186 | 176 | 158 | 127 | 41 | 152 | 188 |
| % of analyzed epochs | 93 | 95 | 97 | 83 | 92 | 68 | 99 | 69 | 92 | 85 | 82 | 86 | 58 | 68 | 53 | 92 |
| Right PAC + SAC | 2 | 2 | 1 | 0 | 1 | 1 | 1 | 1 | 0 | 2 | 0 | 0 | 0 | 0 | 2 | 0 |
| Left PAC + SAC | 0 | 2 | 0 | 2 | 0 | 0 | 0 | 0 | 2 | 2 | 1 | 1 | 0 | 2 | 0 | 0 |

